# Ineffective behavioral rescue despite partial brain Dp427 restoration by AAV9-U7-mediated exon 51 skipping in *mdx52* mice

**DOI:** 10.1101/2025.09.19.677185

**Authors:** Ophélie Vacca, Amel Saoudi, Mathilde Doisy, Xaysongkhame Phongsavanh, Olivier Le Coz, Cathy Nagy, Julia Kuzniar, Cyrille Vaillend, Aurélie Goyenvalle

## Abstract

The *mdx52* mouse model exhibits a common mutation profile associated with brain involvement in Duchenne muscular dystrophy (DMD), characterized by heightened anxiety, fearfulness, and impaired associative fear learning. Deletion of exon 52 disrupts the expression of two dystrophins found in the brain (Dp427 and Dp140), and is eligible for therapeutic exon-skipping strategies. We previously demonstrated that a single intracerebroventricular administration of an antisense oligonucleotide (ASO) targeting exon 51 of the *Dmd* gene could restore 5% to 15% of Dp427 expression. This treatment reduced anxiety and unconditioned fear in *mdx52* mice, improved fear conditioning acquisition, and partially improved fear memory tested 24 hours later. To improve the restoration of Dp427 and induce a long-lasting therapeutic effect, we employed a vectorized approach using an AAV-U7snRNA vector to deliver antisense sequences to the brains of *mdx52* mice. We evaluated two AAV serotypes known for their brain transduction efficiency (AAV9 and RH10) and two delivery routes, intracisterna magna and intracerebroventricular (ICV) injections, to maximize brain targeting. Based on GFP expression data, we selected the AAV9 capsid and a bilateral ICV delivery route. Using this approach, we demonstrated that ICV administration of AAV9-U7-Ex51M induced exon 51 skipping and restored Dp427 expression in the brains of adult *mdx52* mice, though with significant variability among individuals. While a few mice showed high Dp427 expression levels, the average restoration was limited to approximately 6% to 12%. In conclusion, inducing exon skipping in the brains of adult *mdx52* mice using the vectorized AAV9-U7 approach was less effective than synthetic ASO treatment and did not improve the emotional behavior of *mdx52* mice.

## INTRODUCTION

Duchenne muscular dystrophy (DMD) is a severe X-linked neuromuscular disorder primarily characterized by progressive degeneration of skeletal and cardiac muscle due to mutations in the *DMD* gene, which encodes dystrophin proteins^1^. However, the pathological aspect of DMD is not limited to muscle loss; the central nervous system is also significantly affected^2–4^. Indeed, the *DMD* gene gives rise to several brain dystrophins, namely Dp427, Dp140, and Dp71, which are differentially expressed in brain structures are crucial for normal neurodevelopment and synaptic function^5, 6^. Loss of these isoforms has been associated with a range of neurocognitive impairments observed in individuals with DMD, such as intellectual disability, learning difficulties, attention deficit hyperactivity disorder (ADHD), autism spectrum disorder, and emotional disturbances including anxiety and behavioral dysregulation^3, 7^. These neuropsychiatric manifestations can profoundly affect the quality of life of patients and their caregivers, adding a substantial burden to the already complex management of the physical aspects of the disease^8^.

Despite these widespread neurocognitive and emotional issues, most current DMD treatments focus exclusively on improving muscle function and extending mobility and lifespan^9, 10^. However, addressing the brain’s involvement is equally vital. Targeted therapies that can restore or compensate for the missing dystrophin isoforms in the brain have the potential to improve cognitive performance, emotional well-being, and overall quality of life. In this context, providing robust evidence of successful dystrophin restoration within the CNS remains essential, particularly for neuromuscular disorders such as DMD, where the absence of full-length dystrophin also contributes to cognitive and behavioral deficits. Emerging approaches, such as antisense oligonucleotides (ASOs) and viral vector-based gene therapies, offer promising avenues to bypass specific gene mutations and restore dystrophin expression also in the brain.

In recent years, considerable efforts have been made by the biotech industry to develop innovative delivery platforms capable of efficiently transporting ASOs beyond biological barriers. Among these, antibody–oligonucleotide conjugates (AOCs) targeting the transferrin receptor—such as those developed by Avidity Biosciences—have shown promise in enhancing ASO uptake into difficult-to-reach tissues, particularly skeletal muscle^11^. While these strategies are not yet optimized for CNS delivery, recent studies have demonstrated that transferrin receptor-targeted vehicles can enable ASO transport across the blood–brain barrier (BBB) in preclinical models^12^. These advances further underscore the importance of thoroughly investigating and characterizing the therapeutic potential of dystrophin restoration in the brain.

Developing brain-targeted treatments is not only a scientific challenge but also an ethical imperative. These therapies could provide patients with a more holistic benefit, addressing both the physical and cognitive dimensions of the disease. Moreover, by improving neurobehavioral outcomes, these treatments could enhance patients’ ability to engage in education, social activities, and daily living, promoting greater independence and reducing the burden on caregivers.

Given that approximately 13–14% of DMD patients carry mutations eligible to exon 51 skipping, this approach represents the most broadly applicable single-exon skipping strategy currently under clinical investigation^13^. Exon skipping therapies aim to restore the *DMD* reading frame, enabling the production of internally truncated but partially functional dystrophin proteins, similar to those found in Becker muscular dystrophy (BMD)^14, 15^. Importantly, exon 51 skipping allows for the restoration of the full-length Dp427 isoform only, as exon 51 contains the start codon for Dp140^16^, which is consequently lost during the process. Therefore, it is essential to first investigate the specific contribution of Dp427 restoration alone in preclinical models, in order to better anticipate its therapeutic potential in DMD patients eligible for exon 51 skipping. Such studies are crucial to understanding the extent to which Dp427 alone can alleviate central nervous system (CNS) symptoms of the disease and to guide the future development of strategies aiming to co-restore multiple dystrophin isoforms, including Dp140.

We previously explored the potential of tricyclo-DNA antisense oligonucleotides (tcDNA-ASOs) to restore dystrophin expression and alleviate brain-related symptoms in *mdx52* mice, a model of DMD with severe central disorders^17, 18^. A single intracerebroventricular (ICV) administration of tcDNA-ASOs targeting *Dmd* exon 51 achieved partial restoration of Dp427 (5-15%) in brain regions expressing dystrophins, including the hippocampus, cerebellum, and cortex. Behaviorally, this treatment significantly reduced anxiety and unconditioned fear while fully rescuing associative fear learning. However, fear memory tested 24 hours post-conditioning was only partially improved. These findings highlighted the potential for postnatal dystrophin restoration to mitigate emotional and cognitive deficits associated with DMD. However, they also pointed to the need to explore alternative restoration strategies to achieve more comprehensive therapeutic outcomes.

The use of modified U7 small nuclear RNA (U7snRNA) to deliver antisense sequences represents a promising alternative strategy for inducing exon skipping in DMD, an approach that we originally pioneered in the *mdx* mouse model to rescue Dp427 in muscles^19^ or locally in brain^20, 21^. We demonstrated that embedding antisense sequences into U7snRNA and systemically delivering them via adeno-associated virus (AAV) vectors enables efficient and sustained exon skipping, leading to the long-term restoration of dystrophin expression in muscles ^21–26^. AAV vectors offer several advantages, including broad tissue tropism, long-term gene expression, and a favorable safety profile, making them appropriate tools to target muscle tissues after systemic delivery, but also suitable for CNS applications^27^. Notably, AAV-based gene therapies have already been employed successfully in clinical settings for CNS disorders by stereotaxic local delivery, such as spinal muscular atrophy and aromatic L-amino acid decarboxylase deficiency, highlighting their therapeutic potential^28^. Given these considerations, we chose to investigate ICV delivery of AAV-U7snRNA vectors to assess their efficacy in restoring dystrophin expression within the brain.

With the aim to increase Dp427 re-expression and improve the therapeutic potential of exon skipping in the brain, we developed an AAV-U7 vector specifically designed to mediate mouse *Dmd* exon 51 skipping in adult *mdx52*. We first determined that AAV9, delivered via local ICV injection, provided the most efficient brain-wide transduction compared with AAV-RH10. Using this strategy, we showed that the AAV9-U7-Ex51M vector led to high levels of Dp427 restoration in several brain regions. However, the distribution and expression levels of Dp427 protein following AAV-U7 treatment remained heterogeneous and varied substantially between animals. Moreover, we did not observe significant improvements in behavior, even when correlating dystrophin levels with emotional parameters.

## RESULTS

### Selection of capsid and administration route for optimal brain transduction

To determine optimal viral transduction conditions, we first compared AAV9 and RH10 serotypes^29, 30^ expressing GFP after bilateral intracerebroventricular (ICV) injection in *mdx52* mice since we previously demonstrated that it was the best delivery route to achieve broad targeting in the brain of *mdx52* mice^18^. Four weeks post-injection of AAV9-CAG-GFP or RH10-CAG-GFP, both serotypes exhibited similar widespread transduction throughout the forebrain, midbrain, and hindbrain **(Figure 1 A-D)**. Stronger GFP expression was observed in the hippocampus, cortex, thalamus, striatum, and cerebellum. In an attempt to increase transduction in the cerebellum, we also evaluated *intracisterna magna* (ICM) injection compared to bilateral ICV injection, as well as a combination of ICM and bilateral ICV. The ICM route resulted in low transduction efficiency, and combining ICM with bilateral ICV did not enhance the transduction pattern **(Figure 1 E-H)**. Consequently, we proceeded with the standard bilateral ICV route, which demonstrated the highest efficiency for delivery.

**Figure 1:**
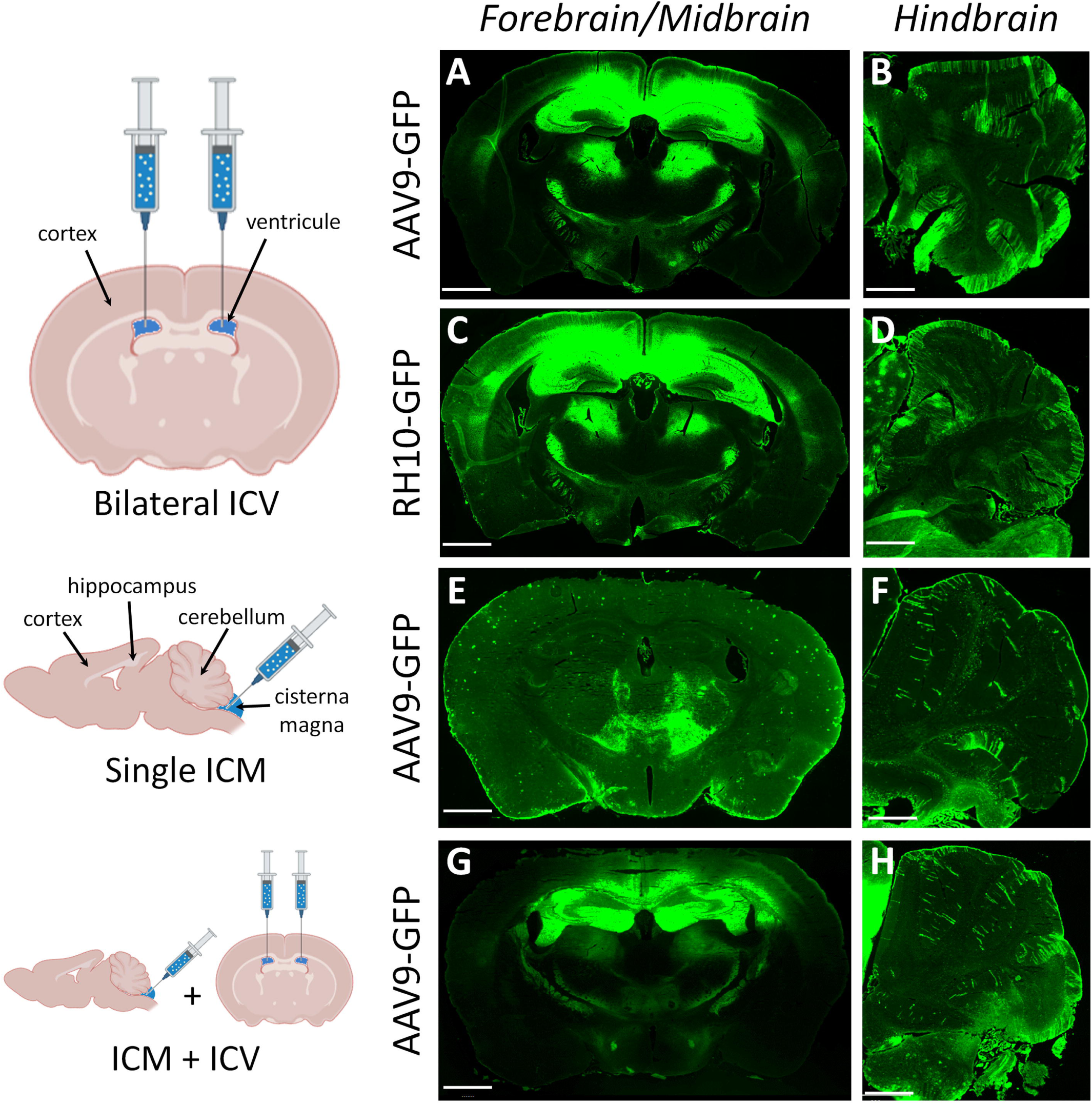
GFP expression patterns in *mdx52* mice following AAV-mediated delivery with various injection strategies. The left panels illustrate the injection methods. **(A-B):** GFP expression 4 weeks after bilateral intracerebroventricular (ICV) injection of AAV9-CAG-GFP. (**C-D):** GFP expression 4 weeks after bilateral ICV injection of RH10-CAG-GFP. (**E-F):** GFP expression 4 weeks after a single intracisterna magna (ICM) injection of AAV9-CAG-GFP. (**G-H):** GFP expression 4 weeks after a combined single ICM and bilateral ICV injection of AAV9-CAG-GFP. Representative images are shown for each treatment group (*n* = 3 mice per group). Scale bar: 1 mm.

At the subcellular level (**Figure 2**), double labeling of GFP with red NeuroTrace staining of Nissl bodies and GFAP revealed transduction of neurons and glial cells in hippocampus, cortex, BLA and cerebellar lobules, for both AAV9 and RH10 serotypes. The majority of transduced cells were neurons, which aligns with the biological context of the study, as Dp427 is predominantly expressed in neurons^31, 32^. Glial labeling was less represented, but included astrocytes in hippocampus, cortex and BLA, and radial Bergmann glia in the molecular layer of cerebellar lobules^33^. AAV9 was selected for subsequent experiments due to its wider use in brain-targeting studies^30^.

**Figure 2:**
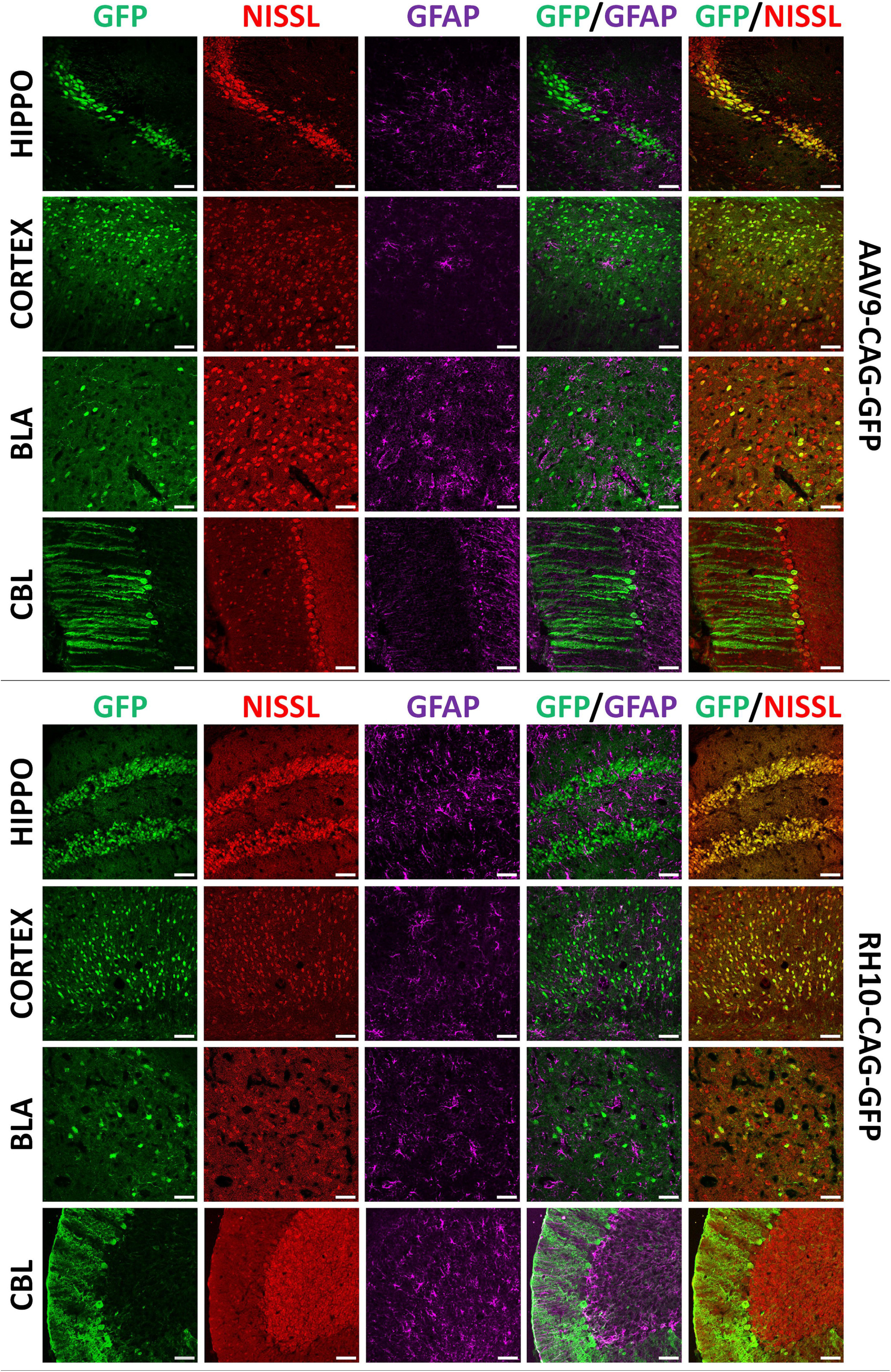
GFP expression patterns in *mdx52* mice following bilateral ICV injections of AAV9 or RH10 serotypes. **(Top):** Confocal images of the hippocampus (HIP), cortex (CX), basolateral amygdala (BLA), and cerebellum (CBL) 4 weeks after bilateral ICV injection of AAV9-CAG-GFP. Sections were labeled with anti-GFP antibody (GFP in green), Nissl marker (nuclei in red), and anti-GFAP antibody (astrocytes in magenta). **(Bottom):** Confocal images of the same regions (HIP, CX, BLA, CBL) 4 weeks after bilateral ICV injection of RH10-CAG-GFP, labeled with the same markers. Representative images are shown for each treatment group (*n* = 3 mice per group). Scale bar: 50 µm.

### Dp427 restoration 9 weeks after bilateral ICV of AAV9-U7-Ex51M

Different U7snRNA constructs were previously engineered to target the human *DMD* exon 51 and were shown to induce efficient skipping of human *DMD* exon 51 both *in vitro* and *in vivo* following intramuscular injection into the tibialis anterior (TA) muscle of a humanized DMD (hDMD) mouse model^25^. Unlike the hDMD mouse model, which carries the entire human *DMD* gene^34, 35^, the *mdx52* mouse model was engineered directly on the murine *Dmd* gene^36^. The antisense sequences of the U7snRNA constructs, originally designed to specifically target human *DMD* exon 51, exhibit a few mismatches with the corresponding mouse exon 51. To address this, we selected the U7 construct that most effectively skips the murine exon 51 while presenting the fewest mismatches. This construct named U7-ex51-long1, contains a single long antisense sequence targeting the region spanning positions +59 to +103 of exon 51^26^ (**Table 1**), that presents two mismatches out of 45 bases (95.5% homology) to the mouse exon 51. We thus corrected these two mismatches via site-directed mutagenesis to achieve 100% homology and maximize exon 51 skipping efficiency in the *mdx52* model (**Table 1**).

**Table 1:**
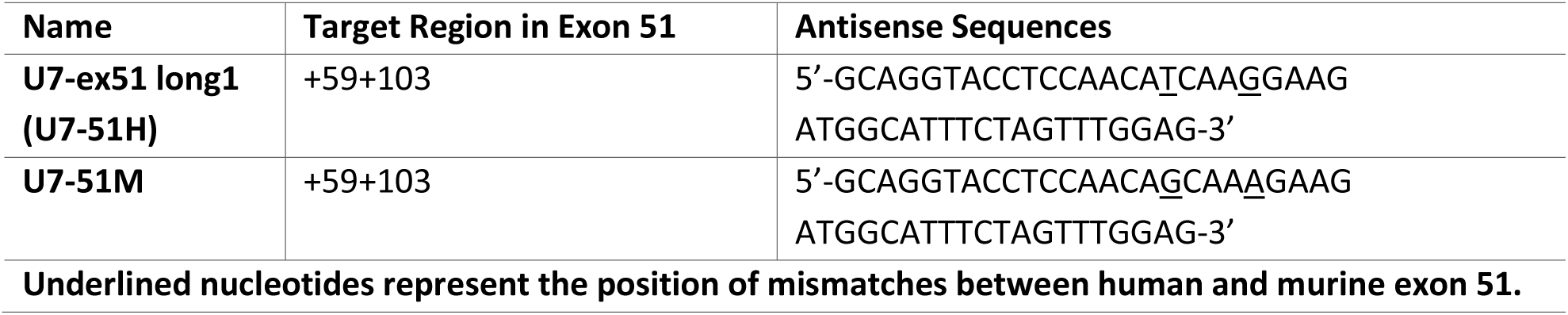
Antisense Sequences Inserted into U7snRNA Constructs Targeting the Human *DMD* Exon 51 and Murine *Dmd* Exon 51.

This optimized U7 construct, named U7-Ex51M (**Figure 3A**), was subsequently cloned into a self-complementary (sc) AAV plasmid and packaged into an AAV9 capsid (**Figure 3B**) for *in vivo* evaluation through intramuscular injection into the TA muscle of *mdx52* mice. The results demonstrated significantly improved exon 51 skipping with the murine U7-EX51M compared to the human U7-ex51-long1 construct three weeks post-injection (**Figure S1**). Based on these findings, we selected the scAAV9-U7-EX51M vector for further *in vivo* evaluations in the *mdx52* mouse model.

**Figure 3:**
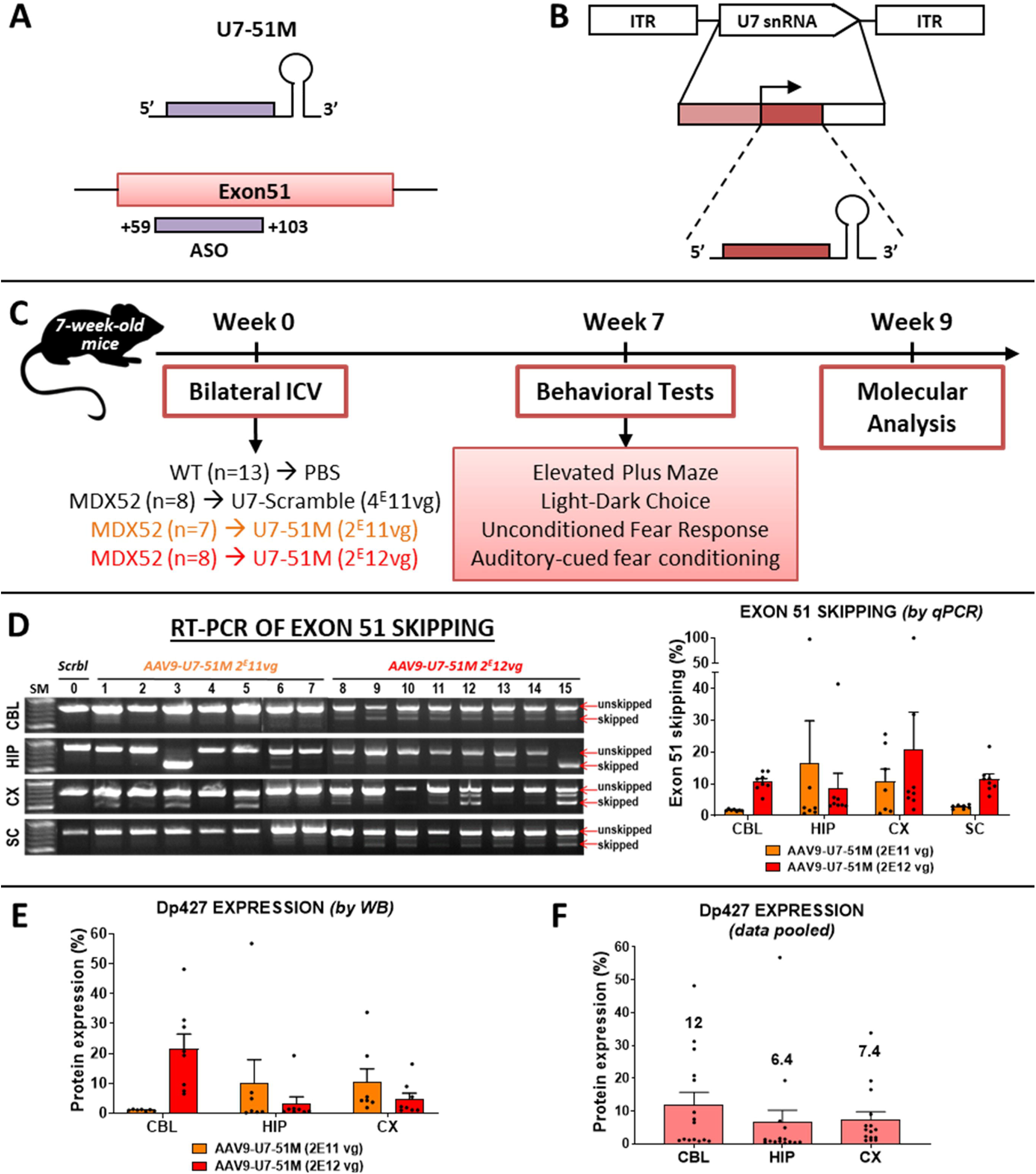
Dp427 restoration 9 weeks after bilateral ICV injection of AAV9-U7-51M. **(A):** Schematic representation of the U7snRNA and its target site on dystrophin pre-mRNA. **(B):** Structure of the AAV vector encoding the U7snRNA cassette. The cassette is flanked by inverted terminal repeats (ITRs) and includes an engineered U7snRNA sequence (gray box) with an antisense region, driven by its natural U7 promoter (hatched box) and 3′ downstream elements (white box). **(C):** Overview of the experimental procedure. **(D): Left panel:** RT-PCR gel showing exon 51 skipping 9 weeks post-injection. SM: DNA ladder; Mouse 0: *mdx52* mouse injected with U7-Scramble (negative control); Mice 1 - 7: *mdx52* mice treated with a low dose of AAV9-U7-51M (1E11 vg); Mice 8 - 15: *mdx52* mice treated with a high dose of AAV9-U7-51M (1E12 vg). **Right panel:** Quantification of exon 51 skipping via RT-qPCR, comparing low- and high-dose treatments (*n*=7 for low dose in orange and 8 for high dose in red). **(E):** Quantification of Dp427 protein restoration by western blot after AAV9-U7-51M injection at low and high doses expressed in a percentage of the WT (*n*=7 for low dose in orange and 8 for high dose in red). Statistical analysis (two-way ANOVA) did not reveal a dose-dependent effect due to high interindividual variability. **(F):** Combined analysis of Dp427 protein rescue levels irrespective of the injected dose expressed in a percentage of the WT (n=15) in the three brain structures.

Our objective was to restore Dp427 expression in the brains of *mdx52* mice with the scAAV9-U7-EX51M vector, to rescue their emotional phenotype. To this end, we performed bilateral ICV injections to four groups of 7-week-old mice: 13 PBS-treated wild-type mice (WT), 8 *mdx52* control mice treated with scAAV9-U7-scramble, 7 *mdx52* mice treated with scAAV9-U7-EX51M at a low dose of 2E11 vg, and 8 *mdx52* mice treated with scAAV9-U7-EX51M at a high dose of 2E12 vg (**Figure 3C**).

Seven weeks post-injection, the mice were analyzed, starting with two weeks of behavioral testing followed by molecular analysis of brain tissues. When measuring exon-skipping levels by both RT-PCR and RT-qPCR, we found a large interindividual variability, ranging from 0.7% to 99% (**Figure 3D**). When considering group means, exon skipping in the low-dose group ranged from 1.7% in the cerebellum (CBL) to 16% in the hippocampus (HIP), while the high-dose group ranged from 8% in the hippocampus (HIP) to 20% in the cortex (CX). We next assessed Dp427 restoration by western blot (WB) and observed the same interindividual variability, ranging from 0.2% to 56.8%. In terms of group means, average protein restoration ranged from 1.08% in the CBL to 10.38% in the CX for the low-dose group, and from 3.25% in the HIP to 21.51% in the CBL for the high-dose group (**Figures 3E and S2**). Despite the presence of a significant number of skipped mRNAs (between 1.4% and 21.69%), we demonstrated in a previous study that we could not detect any protein restoration in the cervical region of the spinal cord^18^. This lack of restoration may be due to the very low levels of Dp427 typically expressed in this structure. For this reason, we did not investigate the restoration of Dp427 in the spinal cord. A two-way ANOVA analysis of data presented in figure 3E revealed no significant differences between the two groups, indicating the absence of a dose-dependent effect, consistent with differences observed in the viral genome distribution in the different brain regions (**Figure S3**). Therefore, data from the two groups were pooled for subsequent analyses (**Figure 3F**). Overall, these results demonstrate an average restoration of Dp427 of 6.4% in the hippocampus (HIP), 7.4% in the cortex (CX), and 12% in the cerebellum (CBL) in *mdx52* mice. It is also important to note that the restoration of protein is lower than the level of skipped transcripts, reflecting a discrepancy previously reported^26^ and attributed to the 5’-3’ transcript imbalance highlighted by prior research^37–39^.

### Partial restoration of Dp427 expression in the brain of AAV9-U7-51M–treated *mdx52* mice

In order to evaluate more precisely the localization of the restored Dp427 detected by WB, we performed immunofluorescence analyses in various brain regions of *mdx52* mice nine weeks after ICV injection of either AAV9-U7-51M or a scrambled control vector (**Figure 4, left**). We used the pan-dystrophin antibody H4, which detects all brain-expressed dystrophin isoforms: Dp427, Dp140, and Dp71. In WT mice, there was a punctate staining that is expected to mainly reflect synaptic expression of Dp427, and a typical staining along blood vessels reflecting perivascular expression of Dp71. In control *mdx52* mice treated with AAV9-U7-Scramble (named MDX52 on the figures), only Dp71 was detected in the examined brain regions, primarily around blood vessels and capillaries (indicated by white stars), resulting in a markedly reduced overall fluorescent signal, consistent with the known absence of full-length Dp427 and Dp140 in these animals. A similar staining was observed in the treated mice, for which we did not detect Dp427 rescue by WB, confirming the absence of protein rescue in some treated animals. In contrast, mice treated with AAV9-U7-51M for which we had detected Dp427 rescue by WB, exhibited partial restoration of dystrophin immunoreactivity, characterized by the typical punctate synaptic staining, in the stratum pyramidale (SP) and proximal stratum radiatum (SR) of the CA1 hippocampal region, as well as in the cortex, basolateral amygdala (BLA), and both the Purkinje cell and molecular layers of the cerebellum. Quantification of the fluorescent area in these mice revealed a significant increase in dystrophin signal (expressed as % of WT levels) in the hippocampus (71%, *p* < 0.0001), cortex (53%, *p* < 0.0001), BLA (30%, *p* = 0.0133), and cerebellum (54%, *p* < 0.0001) as compared to scramble-treated controls (Kruskal–Wallis test followed by Dunn’s post-hoc test) (**Figure 4, right**). Furthermore, dystrophin expression levels in these AAV-treated mice were no longer significantly different from those in WT mice, except in the BLA, where they remained significantly lower (**p = 0.0078). These findings demonstrate that when successful, the AAV9-U7-51M has the potential to locally rescue significant levels of Dp427 in the brain of mdx52 mice; however, this restoration remains partial and spatially heterogeneous throughout the brain.

**Figure 4.**
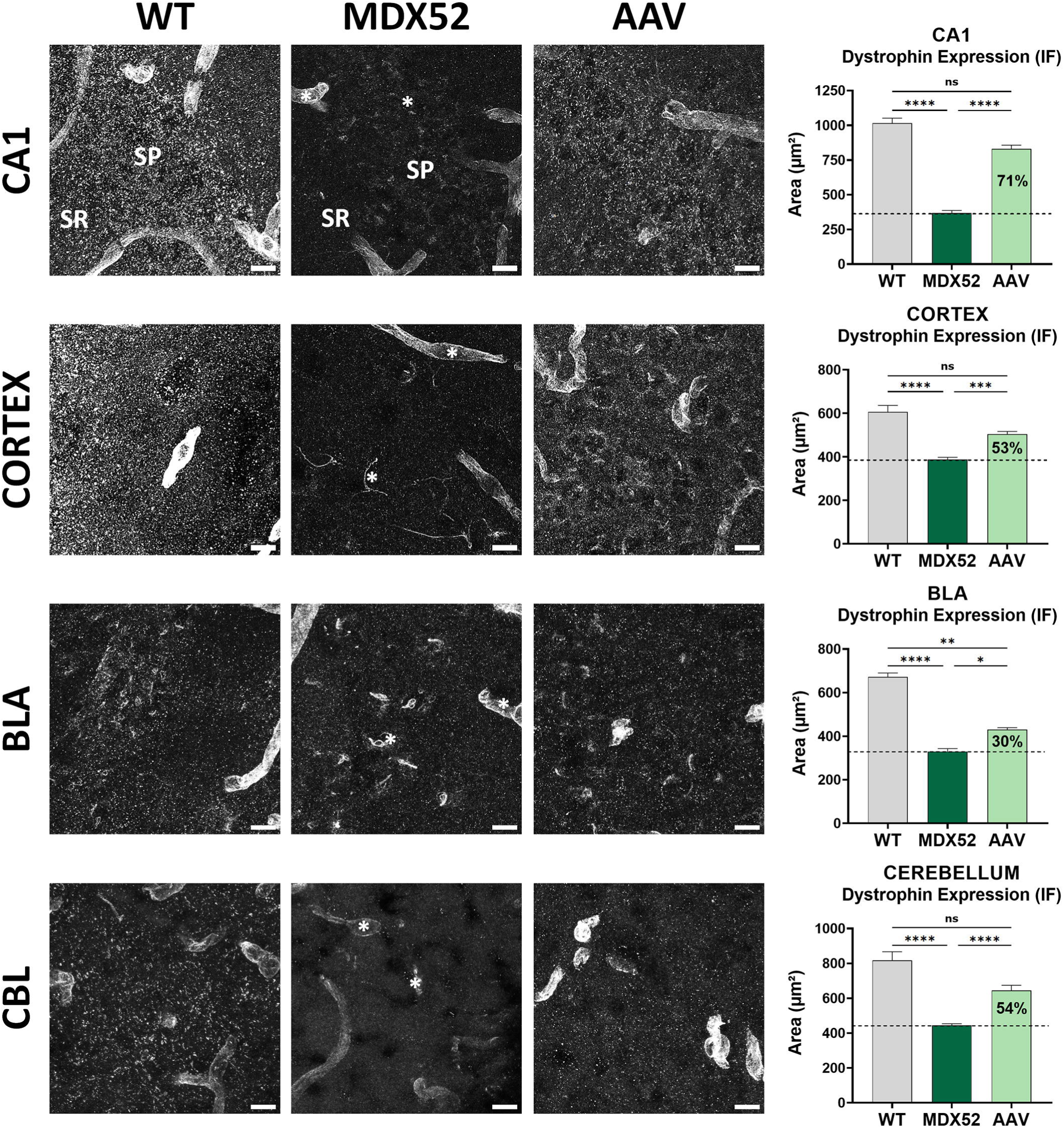
Localization and quantification of Dp427 re-expression. **Left panels:** Representative confocal images of dystrophin immunofluorescence using the pan-dystrophin antibody H4 in hippocampus (CA1 region), cortex, basolateral amygdala (BLA), and cerebellum, nine weeks after intracerebroventricular (ICV) injection of AAV9-U7-Scramble (MDX52) or AAV9-U7-51M (AAV), compared to wild-type (WT) mice. In *mdx52* mice, staining corresponds to Dp71 expression around blood vessels and capillaries (white stars). Restored Dp427 was identified as punctate post-synaptic labelling in the stratum pyramidale (SP) and stratum radiatum (SR) of the hippocampus, the molecular and Purkinje cell layers of the cerebellum, the cortex, and the BLA. Scale bars = 10 µm. **Right panel:** Quantification of dystrophin-positive fluorescent area (µm²) in each brain region. WT values represent the combined expression of Dp427 and Dp140, while in treated *mdx52* mice, signal above MDX52 levels reflects restored Dp427. Data are shown as mean ± SEM; percentages in bars indicate expression relative to WT. Statistical significance was determined using Kruskal–Wallis test followed by Dunn’s multiple comparisons; *p < 0.05, **p < 0.01, ***p < 0.001, ****p < 0.0001. Sample sizes: n = 4–8 mice per group.

### Impact of Partial Dp427 Restoration on Neurobehavioral outcomes in *mdx52* Mice Following AAV9-U7-Ex51M ICV Administration

The behavioral study was conducted on all groups: WT controls, untreated and treated *mdx52* mice, starting with an assessment of anxiety in the elevated plus maze test (**Figure 5A**). In this test, the percent of time spent in the center and in the open arms of the maze, the most anxiogenic areas, was measured. As previously shown, *mdx52* mice spent significantly less time in the center and in the open arms (*p* < 0.0001)^17^. Mice treated with AAV-U7-EX51M also spent significantly less time in the center and in the open arms (*p* < 0.0001), displaying a behavior similar to that of *mdx52* control mice (**Figure 5B**).

**Figure 5:**
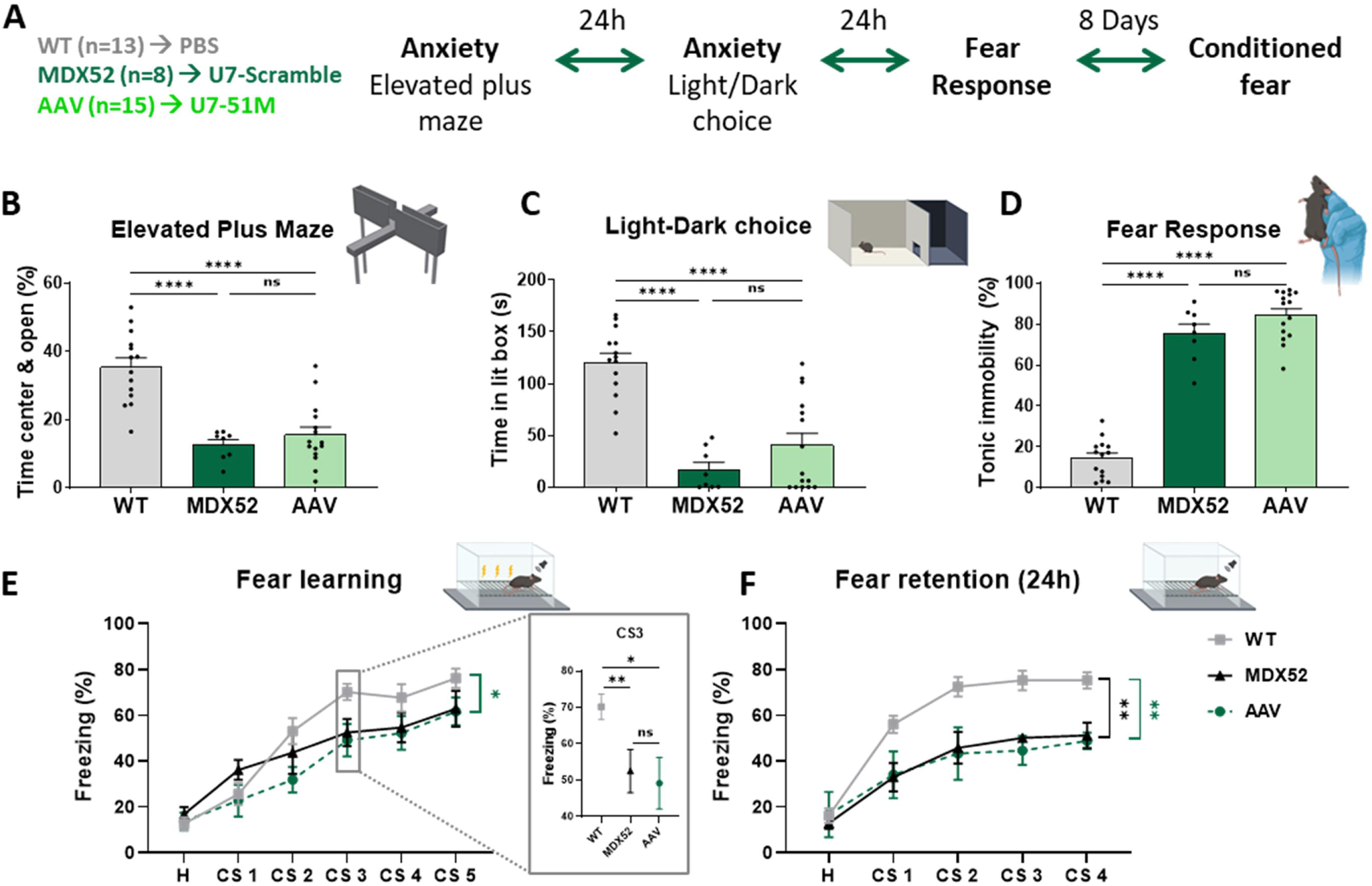
Impact of Partial Dp427 Restoration on Emotional Reactivity and Emotional Learning in *mdx52* Mice Following AAV9-U7-Ex51M ICV Administration. (A) Behavioral testing, initiated seven weeks post-ICV, assessed emotional reactivity in treated (*n*=15 AAV9-U7-51M) and control mice (*n*=13 WT PBS, *n*=8 *mdx52* AAV9-U7-Scramble). Tests included the elevated plus maze and light/dark choice tests, followed by unconditioned fear response measurement (24-hour intervals between tests), and concluded with auditory-cued fear conditioning eight days later. **(B)** Elevated plus maze: Time spent in the center and open arms (percentage of test duration). \*\*\*\**p* < 0.0001, one-way ANOVA with Holm-Sidak post-hoc test. **(C)** Light/dark choice test: Time spent (s) in the lit compartment. \*\*\*\**p* < 0.0001, one-way ANOVA with Holm-Sidak post-hoc test. **(D)** Unconditioned fear response: Percentage of time spent in tonic immobility during a 5-minute recording following 15 seconds of restraint. \*\*\*\**p* < 0.0001, one-way ANOVA with Holm-Sidak post-hoc test. **(E)** Auditory-cued fear conditioning: Percentage of time spent freezing during conditioned stimulus (CS) (30-second, 80 dB tone) presentation. Conditioning comprised five coupled stimuli (tone followed by foot shock), and **(F)** retention was assessed 24 hours later in a novel context (tone only). \**p* < 0.05, \*\**p* < 0.01, one-way ANOVA with Holm-Sidak post-hoc test. The insert corresponds to the third conditioned stimulus (CS3) during the fear learning session analyzed by Mann–Whitney. \**p* < 0.05, \*\**p* < 0.01. All data are represented as means ± SEM.

Twenty-four hours later, the mice underwent the light-dark choice test, which evaluates anxiety triggered by intense light. As established in a previous study, *mdx52* control mice spent significantly less time in the illuminated part of the box compared to WT mice (*p* < 0.0001)^17^. Similarly, mice treated with AAV-U7-EX51M spent significantly less time in the illuminated area than WT mice (*p* < 0.0001), exhibiting behavior akin to that of *mdx52* control mice (**Figure 5C**).

Twenty-four hours after this, the mice were tested for their unconditioned fear response to an acute stress induced by a 15-second restraint. *Mdx52* control mice display a strong phenotype, with a significantly larger percentage of tonic immobility compared to WT mice (*p* < 0.0001)^17^. *Mdx52* treated mice did not show any reduction in this tonic immobility (*p* < 0.0001) (**Figure 5D**).

Eight days later during which the mice were handled, the three groups were tested in an associative fear conditioning task based on the coupling of auditory cues (CS) and aversive stimuli (US). This test consists of an acquisition session, during which the mouse learns to associates a tone that predicts the delivery of an electric foot shock. The *mdx52* mice exhibited no fear learning deficit during acquisition (p = 0.5169 with WT), except during delivery of the third conditioned stimulus (*p* = 0.0060, analyzed by Mann-Whitney). The *mdx52* mice treated with AAV9-U7-EX51M did not differ from the other experimental groups (p = 0.6374 with untreated *mdx52*, *p* = 0.0469 with the WT). Moreover, *mdx52* presented a fear memory deficit (*p* = 0.0013)^17^, which was not improved in the AAV9-U7-EX51M – treated mice (*p* = 0.9971 with *mdx52*, *p* = 0.0024 with the WT) (**Figure 5E**).

In summary, we did not observe any improvement in the behavioral phenotype in either the anxiety or the fear response or the fear memory tests.

### Complementary analysis of the behavioral outcomes taking in account mice re-expressing a minimum of 4% of Dp427 in brain

Due to the substantial inter-individual variability in Dp427 restoration levels across treated mice, a complementary analysis was conducted to focus specifically on a subset of animals showing a consistent average restoration of at least 4% Dp427 across the three analyzed brain regions: cerebellum (CBL), hippocampus (HIP), and cortex (CX). This threshold was selected based on our previous findings with ASO treatment, which indicated that behavioral improvements were observed at this range of dystrophin restoration.^18^ A total of 9 mice met this criterion and were included in the analysis. These animals are indicated with an asterisk in Table 2. This approach aimed to determine whether a minimum threshold of 4% Dp427 expression was sufficient to rescue specific behavioral phenotypes.

**Table 2:**
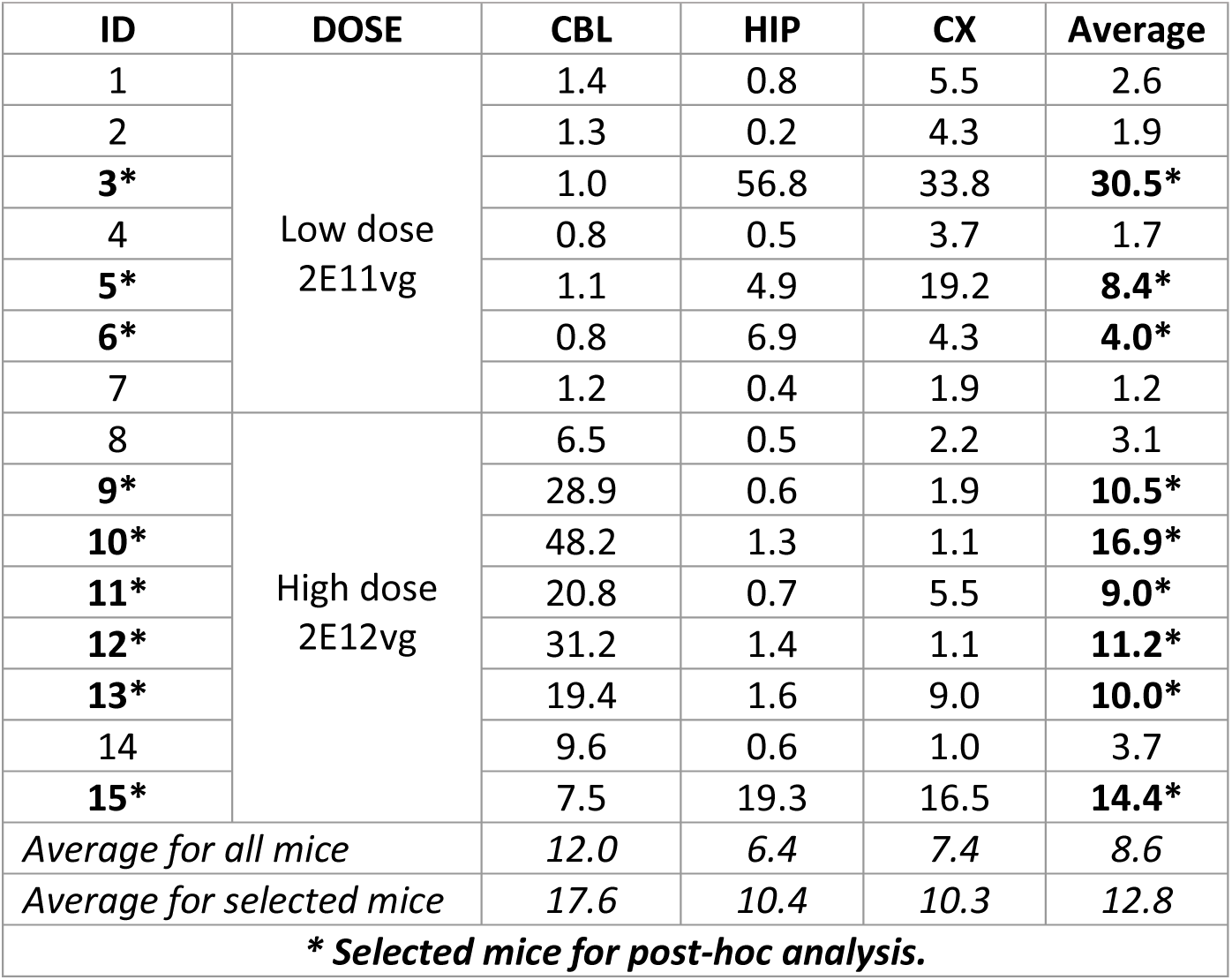
Selected mice with an average of at least 4% Dp427 expression across all brain regions (CBL, HIP, CX) for post-hoc analysis.

In average, the selected mice exhibited Dp427 rescue levels of 17.2% in the CBL, 10.4% in the HIP, and 10.3% in the CX (**Figure 6A**).

**Figure 6:**
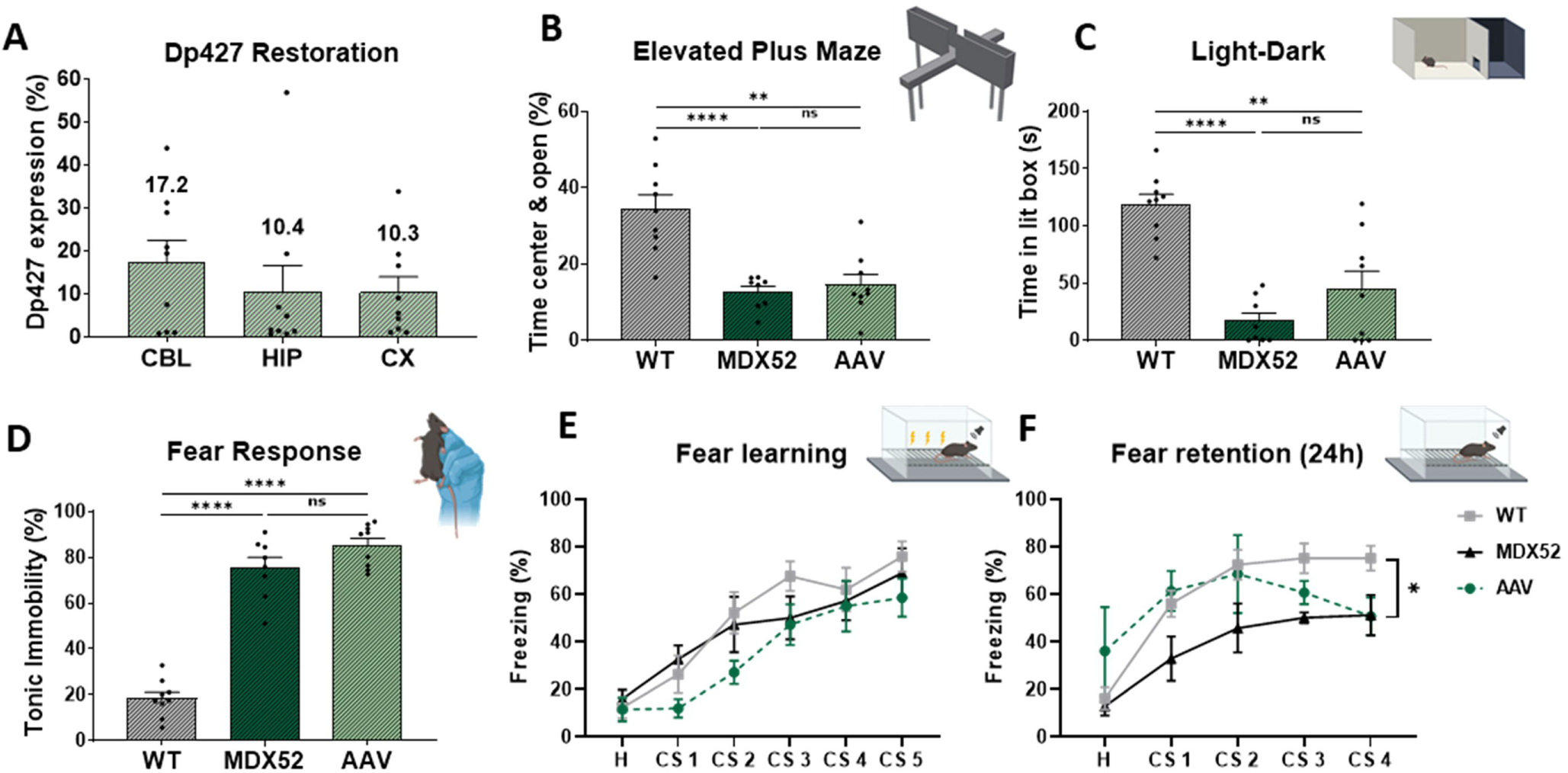
Behavioral Analysis in a selected subset of mice. Mice exhibiting a minimum of 4% Dp427 re-expression across all brain regions (CBL, HIP, CX) were selected for further analysis (n=9). **(A)** Percentage of Dp427 restoration in the selected mice for CBL, HIP, and CX. The evaluation of behavioral improvements was conducted through various tests: the elevated plus maze **(B)**, the light-dark choice test **(C)**, the unconditioned fear response test **(D)**, and the auditory-cued fear conditioning test **(E, F)** using the same cohort of post-hoc selected mice (n=9). Data from panels B, C, and D are presented as mean ± SEM and analyzed using Mann-Whitney test; **p < 0.01, ****p < 0.0001. Data for panels E and F are expressed as mean ± SEM and analyzed with two-way ANOVA and Sidak post-hoc test; *p < 0.05.

In the behavioral analysis of this subset of mice, we observed a very slight improvement in both anxiety-related behavioral tests—the elevated plus maze (EPM) and light–dark box (LD)—as AAV-treated mice showed a slightly reduced difference compared to WT mice. However, they remained not significantly different from *mdx52* control mice, and still significantly different from WT mice (**Figure 6B–C**). No significant improvement was observed in unconditioned fear responses or in fear memory (**Figure 6D–F**). We also investigated whether Dp427 restoration levels correlated with anxiety- or fear-related behaviors, considering both the entire group of treated mice and the selected subset. No significant correlations were found between Dp427 levels and behavioral outcomes whether considering all treated mice or the mice chosen for the post-hoc analysis (**Figure S3**).

## DISCUSSION

In this study, we first assessed the tropism of two AAV serotypes, AAV9 and AAVrh10, in the brain of *mdx52* mice, using three different delivery routes: intracerebroventricular (ICV), intracisterna magna (ICM), and a combination of both (ICV + ICM). Our results indicate that AAV9 and AAVrh10 exhibit broadly similar transduction patterns across brain regions, in agreement with previous reports showing overlapping CNS tropism for these two serotypes following ICV administration^30, 40, 41^. Furthermore, we found that ICV delivery alone was as efficient as the combined ICV + ICM approach, suggesting that ICM does not provide additional therapeutic benefit in terms of brain-wide transduction.

We then aimed to restore widespread Dp427 expression in the *mdx52* mouse brain by bilateral ICV injection of two different doses of AAV9-U7-Ex51M. Nine weeks post-treatment, we observed no clear dose-dependent effect and detected high inter-individual variability in exon 51 skipping levels which was in line with the viral genome quantification by brain region (**Figure S4**). Of note, we never observed such variability in our previous studies using bilateral ICV injection of ASOs, confirming that this is not related to the technical procedure, which is well established and routinely performed in our laboratory^18, 42–44^. Such variability is consistent with the variable penetrance of ICV-mediated gene delivery previously described in the literature that can be attributed to factors such as cerebrospinal fluid (CSF) dynamics and individual anatomical differences^45, 46^.

In line with exon skipping results, we also observed a marked inter-individual variability in Dp427 protein restoration, ranging from 0.2% to 56.8% (Table 2), as well as intra-brain region variability, with some regions showing high levels of rescue while others appeared not restored at all. Indeed, mice exhibiting high levels of Dp427 rescue in one brain region can display very low levels in another (e.g. mouse 10 with the highest rescue in CBL, showed extremely low rescue in HIP and CX; in contrast, mouse 3 with the highest rescue in HIP, showed very low rescue in CBL). Notably, samples with the highest exon skipping levels also exhibited the greatest protein restoration, and the same trend was observed for those with the lowest levels. Despite this particularly high variability, it is interesting to note that AAV-mediated delivery, when successful, resulted in higher peak Dp427 expression restoration compared to our previous work with tcDNA, which did not exceed 25%^18^. This confirms that AAV technology has the potential to induce robust dystrophin restoration; however, further optimization of vector biodistribution remains necessary to enhance therapeutic efficacy.

Given this large inter-individual variability and the lack of a clear dose-response relationship, we decided to pool data across both treatment groups for behavioral analysis. Despite this approach, we were unable to detect significant behavioral improvements in the full cohort. We hypothesized that animals exhibiting very low levels of Dp427 restoration may have masked potential treatment effects. To address this, we conducted a complementary analysis focusing on a subset of nine mice that exhibited at least 4% Dp427 expression across all brain regions. Within this subgroup, we observed very modest and non-significant improvements in anxiety-related behaviors and fear memory.

We performed correlation analyses between Dp427 restoration levels and behavioral outcomes considering all mice or only in the selected subset of mice, but no correlation could be established (**Figure S2**). We also tried to correlate behavior with restoration levels by region, but this also did not correlate (data not shown). Notably, this analysis did not exclude animals with 0% Dp427 expression in individual brain areas, which may have limited the observed behavioral gains. In contrast, our previous study using tcDNA-Ex51 achieved more consistent Dp427 restoration across brain regions, albeit at relatively low levels (5–15%), and this was sufficient to partially rescue emotional phenotypes in *mdx52* mice. These observations suggest that a homogeneous rescue of Dp427 across brain regions is likely required to achieve meaningful therapeutic outcomes even with quite low levels.

In recent years, several clinical trials have demonstrated the therapeutic potential of AAV-based gene therapies for monogenic neurological disorders^47, 48^. Strategies employing AAV9 or AAVrh10 capsids have been developed to deliver genetic material directly to the CNS, often through intrathecal or intracerebroventricular administration routes^49^. These approaches aim to enhance vector distribution across brain regions while minimizing systemic exposure and associated immune responses. Promising outcomes have been reported in conditions such as GM1 gangliosidosis^50^, Canavan disease^51^, and Dravet syndrome^52^. These studies report improvements in motor function, reduction of pathological biomarkers, or partial restoration of gene function. Nonetheless, significant challenges remain, including inter-individual variability in transgene expression, immune responses against the viral capsid, and potential toxicity linked to excessive or ectopic gene expression. These limitations highlight the need for further refinement of vector design, delivery parameters, and gene regulation strategies to ensure safety and efficacy in future CNS-directed therapies.

To improve the distribution of recombinant AAV vectors in the brain, alternative strategies would be worth investigating in future experiments. Among them, neonatal ICV administration would be of particular interest since it has been shown to promote broader vector diffusion and more homogeneous transduction in the CNS compared to adult ICV delivery, due to the immature brain parenchyma and enhanced CSF dynamics in neonates^53, 54^. In addition, the use of alternative bioengineered capsids such as AAV-DJ, which have demonstrated improved tissue penetrance and broader diffusion following local injection in adult and neonatal brains would be worth exploring^55–58^. Furthermore, wider brain distribution may be achieved via systemic delivery approaches using capsids that efficiently cross the blood-brain barrier (BBB), such as AAV-PHP.eB, which enables widespread CNS transduction after intravenous injection in rodents^59^. However, while systemic administration can extend vector coverage, it may limit our ability to precisely assess the role of brain dystrophins in behavioral phenotypes. Indeed, peripheral restoration of dystrophin has been shown to influence behavioral outcomes, potentially via neuromuscular feedback or systemic physiological effects, complicating the interpretation of centrally mediated functions^60^. To further dissect the contribution of central dystrophins to behavior, it would be useful to restore both Dp427 and Dp140 isoforms, which are missing in the *mdx52* mouse model. Combined restoration of these isoforms may provide novel insights into their complementary or synergistic contributions to emotional and cognitive functions, and help define isoform-specific therapeutic strategies.

In conclusion, our study demonstrates that AAV9 and RH10 vectors exhibit relatively similar tropism in the CNS of *mdx52* mice when administered via direct ventricular injection. Among the tested delivery routes, bilateral ICV injection appears to be the most effective method, outperforming ICM alone and achieving outcomes comparable to the combined ICM + ICV approach on its own. Despite achieving partial restoration of Dp427 in several brain regions, our results highlight the current limitations of AAV-U7–mediated exon skipping for central nervous system delivery. While AAV9 delivery to the adult brain is effective to some extent, its distribution remained heterogeneous across animals, and this variability likely contributed to the absence of consistent behavioral recovery compared to ASO-based therapies. These findings suggest that, while the approach is feasible, it does not yet provide sufficient efficiency to support functional benefit. Alternative strategies, such as AAV-mediated delivery of specific isoforms, are already under investigation in our laboratory and may help clarify the respective contributions of different dystrophin isoforms to neuronal function. Importantly, we believe that sharing these negative but rigorously collected data is essential for the neuromuscular and gene therapy communities. As emphasized by others, reporting both successes and limitations is critical to guide the field toward more effective and translationally relevant therapeutic approaches.

## MATERIALS AND METHODS

### Animals

The *mdx52* mouse model of X-linked muscular dystrophy, characterized by a deletion of exon 52 in the *Dmd* gene, was developed by Dr. Katsuki Motoya’s team. This model was created by replacing exon 52 with a neomycin resistance gene, resulting in the absence of Dp427, Dp260, and Dp140 dystrophins, while maintaining the expression of Dp116 in peripheral nerves and Dp71 in the brain and retina. The strain was backcrossed with C57BL/6J mice for over eight generations to ensure genetic stability. This mouse line was generously shared by Prof. Toshikuni Sasaoka (Department of Comparative & Experimental Medicine/Brain Research Institute, Niigata University, Japan). Breeding pairs were provided to our laboratory by Dr. Jun Tanihata and Dr. Shin’ichi Takeda (National Center of Neurology and Psychiatry, Tokyo, Japan). In our facility at UVSQ in Montigny-le-Bretonneux (France), heterozygous females were bred with *mdx52* males to produce *mdx52* males and their C57BL/6J wild-type (WT) littermates. Genotyping was performed using PCR analysis of tail DNA. Following treatment, litters were transferred to the animal facility at NeuroPSI (France) for behavioral testing. All animal care and experimental protocols adhered to European regulations (CEE 86/609/EEC and EU Directive 2010/63/EU) as well as French national guidelines (87/848) and were approved by the Paris Center-South Ethics Committee (no. 59).

### Construction of U7snRNA Vectors and AAV Production

U7snRNA constructs targeting the murine *Dmd* exon 51 were engineered based on the previously described U7smOPT-SD23/BP22 backbone^61^ (modified murine U7snRNA gene). The antisense sequence originally targeting human exon 51 was adapted to the murine sequence by site-directed mutagenesis, introducing two specific nucleotide changes (Table 1). The U7-ex51 long1 construct contains a single 45-mer antisense sequence complementary to positions +59 to +103 of murine exon 51 (Table 1).

The resulting U7snRNA fragments were cloned between the inverted terminal repeats (ITRs) of a self-complementary AAV vector (pscAAV, Addgene plasmid #83279; gift from Mark Kay) for subsequent AAV production. We used a self-complementary AAV because scAAV vectors are reported to achieve faster and more robust expression than single-stranded AAV vectors ^62–64^, and the U7 cassette fit within the size limit. AAV9 and RH10-pseudotyped vectors were generated by triple co-transfection of HEK293T cells with the pAAV-U7 plasmid, pXX6 plasmid encoding adenoviral helper functions, and pAAV2/9pITRCO2 plasmid containing the *AAV9* rep and cap genes. Seventy-two hours post-transfection, vector particles were purified from cell lysates using iodixanol gradient ultracentrifugation. Viral genome titers were determined by quantitative real-time PCR as previously described^65^.

### Stereotaxic Surgery and experimental groups

Intracerebroventricular (ICV) injections were conducted on 6- to 7-week-old male *mdx52* and WT littermate mice under deep anesthesia, induced by a single intraperitoneal injection of ketamine (95 mg/kg) and medetomidine (1 mg/kg). A self-complementary adeno-associated virus serotype 9 or RH10 (scAAV9 or a scRH10) vector encoding either the GFP under the control of the CAG promoter (scAAV9-CAG-GFP or scRH10-CAG-GFP) or a modified U7snRNA designed to target dystrophin exon 51 (scAAV9-U7-51M), or phosphate-buffered saline (0.1 mol/L) solutions were bilaterally injected into the lateral brain ventricles (approximately 0.5 mm posterior to bregma, 1 mm lateral, and 2 mm below the dura mater). Each ventricle received an infusion of 5 µL at a rate of 0.5 µL/min, resulting in a total bilateral administration of 2E+11 vg of scAAV9-U7-51M for a low dose group of 7 *mdx52* mice, of 2E+12 vg of scAAV9-U7-51M for a high dose group of 8 *mdx52* mice, of 2+12 vg of scAAV9-U7-Scramble for a control group of 8 *mdx52* mice and of 10 µL of PBS for control group of 13 wild-type (WT) mice. To ensure balanced distribution, the treatment was pseudorandomized within each cage, maintaining comparable allocation within and across litters.

Littermates were housed in groups of two to five per cage under a 12-hour light/dark cycle (lights on at 7:00 a.m.) with unrestricted access to food and water.

### Tissue collection

Following dissection, one hemisphere of the brain was immediately frozen on dry ice for *in situ* analysis, while the other hemisphere was used for the isolation of specific brain regions, including the hippocampus (HIP), cerebellum (CBL), cortex (CX), and the cervical segment of the spinal cord (SC). These tissue samples were snap-frozen in liquid nitrogen for subsequent viral genome biodistribution analysis, qPCR, and western blot experiments.

### Real-Time Quantitative PCR

Total RNA was extracted from dissected brain regions using the TRIzol reagent, following the manufacturer’s protocol (Thermo Fisher Scientific). To assess exon-skipping efficiency via gel electrophoresis, 1 µg of total RNA was subjected to RT-PCR using the Access RT-PCR System (Promega) in a 50 µL reaction volume, with external primers Ex49F (5’-AAACTGAAATAGCAGTTCAAGC-3’) and Ex53R-Aoki (5’-ACCTGTTCGGCTTCTTCCTT-3’). Reverse transcription was performed at 55 °C for 10 minutes, followed by 30 PCR cycles consisting of 95 °C for 30 s, 58 °C for 1 min, and 72 °C for 1 min. PCR products were separated on 1.5% agarose gels. Exon 51 skipping was also quantified by TaqMan real-time PCR as previously described^26^, using specific assays targeting the exon 50-51 junction (assay Mm.PT.58.41685801; forward primer: 5’-CAAAGCAGCCTGACCGT-3’; reverse primer: 5’- TGACAGTTTCCTTAGTAACCACAG-3’; probe: 5’-TGGACTGAGCACTACTGGAGCCT-3’) and the exon 50-53 junction (forward primer: 5’-GCACTACTGGAGCCTTTGAA-3’; reverse primer: 5’- TTCCAGCCATTGTGTTGAATC-3’; probe: 5’-ACAGCTGCAGAACAGGAGACAACA-3’) (Integrated DNA Technology). For each reaction, 150 ng of cDNA was used, and all measurements were performed in triplicate. qPCR was conducted under fast cycling conditions using a Bio-Rad CFX384 Touch Real-Time PCR Detection System. Data analysis was based on the absolute quantification method: the copy numbers of the exon-skipped (exon 50-53) and unskipped (exon 50-51) products were determined using gBlock standards Ex49-54Delta52 and Ex49-54Delta51+52 (Integrated DNA Technology). Exon 51 skipping levels were finally expressed as the percentage of total dystrophin transcripts, calculated as the ratio of skipped copies to the sum of skipped and unskipped copies.

### Western blot

Protein extracts were prepared from brain tissues using RIPA lysis and extraction buffers (Thermo Fisher Scientific), supplemented with SDS to a final concentration of 5% (Bio-Rad, France). Protein concentrations were measured with the BCA Protein Assay Kit (Thermo Fisher Scientific). Samples were denatured at 100 °C for 3 min, and 25 µg of protein per sample was loaded onto NuPAGE 3–8% Tris-acetate gels (Invitrogen), according to the manufacturer’s protocol. Dystrophin expression was analyzed using the anti-dystrophin monoclonal rabbit antibody (ab154168, Abcam, France), while vinculin served as a loading control and was detected with the hVin-1 monoclonal mouse antibody (V9131, Sigma, Saint-Louis, USA). Detection was performed with an IRDye 800CW goat anti-mouse secondary antibody and an IRDye 680CW goat anti-rabbit secondary antibody (Li-Cor, Germany), and bands were visualized using the Odyssey CLx imaging system (Li-Cor). Quantification was carried out with Image Studio software (Li-Cor), normalizing dystrophin signals to vinculin levels. Standard curves were generated for each brain region by mixing lysates from wild-type (WT) and *mdx52* control samples to obtain defined dystrophin percentages (0%, 2.5%, 5%, 10%, and 20% relative to WT tissues).

### Immunochemistry

Coronal and sagittal brain sections (30 µm) were processed for Dp427 staining, tissue sections were first fixed in an acetone/methanol solution (1:1) at –20 °C for 5 min, then permeabilized in PBS containing 0.5% Triton X-100 for 10 min. Slices were subsequently incubated in a blocking solution (10% normal goat serum, 0.1% Triton X-100, 1% BSA, and 1% FAB) for 30 min at room temperature (RT), followed by overnight incubation at 4 °C with a rabbit polyclonal anti-dystrophin antibody (pan-specific H4 antibody, homemade by D. Mornet; 1:1000 dilution in blocking buffer). After washing in PBS, sections were incubated with an Alexa Fluor 488-conjugated secondary antibody (1:400, 1 hour, RT) and counterstained with DAPI. Control sections processed without the primary antibody showed no specific staining. Imaging was performed at equivalent anatomical locations and under identical acquisition settings using a Zeiss LSM 700 laser scanning confocal microscope equipped with a ×60 objective. For each sample, z-stacks of 12 optical sections (1024 × 1024 pixels; 90.19 nm/pixel resolution; 1 µm z-step) were acquired to ensure consistent sampling across animals. Quantification of Dp427 re-expression was conducted in the CA1 region of the hippocampus, the cortex, and the basolateral amygdala (BLA) on coronal sections (bregma –2.0 mm), and in the cerebellum on sagittal sections. For each brain region, multiple sections per mouse were analyzed (3–4 sections), and the mean value per animal was used for statistical analysis. Samples included three untreated control mice and eight treated mice per region. Signal intensity was measured using Fiji software after background subtraction, and expressed relative to control values. This standardized procedure ensured comparability between groups and minimized variability due to acquisition or analysis settings.

### Behavioral Testing

#### Elevated Plus Maze

The apparatus consisted of two closed arms (20 × 8 × 25 cm) with opaque walls and two open arms (20 × 8 cm) without side walls, arranged in a cross shape with a central platform (8 × 8 cm). The maze was elevated 65 cm above the floor. Light intensity was set to 150 lux in the open arms and 30 lux in the closed arms. Each mouse was placed individually at the center of the maze, facing one of the closed arms, and behavior was recorded for 5 min. The number of entries and the time spent in each type of arm were manually scored using event-recorder keys in ANY-Maze software (Stoelting, USA).

#### Light-Dark Box Test

The testing apparatus was composed of a dark chamber (15 × 15 cm, <15 lux) and a brightly lit chamber (40 × 15 cm), connected via a small passage (6 × 6 cm). Illumination in the bright compartment was provided by a lamp positioned at the far end, generating a gradient of light from 50 lux near the doorway to 600 lux at the light source, as described previously^17^. Mice were initially confined to the dark compartment for 10 s before being allowed free access to both compartments for 5 min. Step-through latency, total entries, and time spent in the illuminated chamber were scored manually using event-recorder keys in ANY-Maze software (Stoelting, USA).

#### Restraint-Induced Unconditioned Fear

To elicit acute stress, the mouse was held by the scruff with the tail secured, and gently inverted so that its ventral side faced the experimenter. After 15 sec of restraint, the animal was placed into a clean unfamiliar cage (24 × 19 cm, 12 cm wall height) under low lighting (60 lux), and behavior was recorded for 5 min using ANY-maze software (Stoelting, USA). Freezing behavior, defined as complete immobility aside from respiration, was used as an index of unconditioned fear and expressed as the percentage of total observation time^66^.

#### Auditory-Cued Fear Conditioning

Fear conditioning was performed using the StartFear system (Panlab, Barcelona) following protocols established in prior studies on *mdx* and *mdx52* mice^67, 68^. The conditioning chamber (25 × 25 × 25 cm) featured three opaque walls, a transparent front door, a grid floor linked to a shock generator (unconditioned stimulus, US), and a ceiling-mounted speaker for sound delivery (conditioned stimulus, CS). Animal movement was detected via a high-sensitivity weight transducer and the chamber was enclosed in a ventilated sound-attenuating box (67 × 53 × 55 cm) on an anti-vibration table with background white noise (60 dB). For acquisition, mice were exposed to a 2-minute baseline period, followed by five CS–US pairings (tone: 80 dB, 10 kHz, 30 s; footshock: 0.4 mA, 2 s), with pseudo-random intertrial intervals (60–180 s). On the following day, retention was tested in a novel environment with altered walls and floor. After 2 minutes of habituation, four CS presentations (30 s each) were delivered, separated by variable intervals (60–120 s). Behavioral responses were analyzed using FREEZING software (Panlab) at 50 Hz sampling. Freezing during CS presentations served as a measure of associative memory.

### Statistical analysis

All data are presented as mean ± SEM. Statistical analyses were performed using GraphPad Prism. Two-way ANOVA was used to compare exon 51 skipping and Dp427 restoration between treatment doses (Figures 3D–F and 5E–F), accounting for both dose and inter-individual variability. Due to non-normal distribution of fluorescence data across brain regions, dystrophin expression (Figure 4) was analyzed using the non-parametric Kruskal–Wallis test followed by Dunn’s multiple comparisons test. Behavioral outcomes from the elevated plus maze, light/dark box, and unconditioned fear response (Figures 5B– D) were analyzed using one-way ANOVA with Holm–Sidak’s post hoc test, suitable for multiple group comparisons with parametric data. For targeted comparisons, including post-hoc analysis of selected mice and single timepoints (Figures 5F insert, 6B–D), the Mann–Whitney test was used. Exon skipping efficiencies in Figure S1 were compared using an unpaired t-test, assuming normal distribution of RT-PCR quantification. Correlations between Dp427 levels and behavioral outcomes (Figure S2) were assessed using Pearson’s correlation coefficient with linear regression. Statistical significance was defined as p < 0.05.

## Supporting information

Supplemental informations

## DATA AVAILABILITY STATEMENT

The primary data for this study are available from the authors upon request.

## ACKNOWLEDGMENTS

This work was funded by the European Union’s Horizon 2020 research and innovation program “Brain Involvement iN Dystrophinopathies” under grant agreement No 847826. It was also supported by Centre National de la Recherche Scientifique (CNRS, France), Institut National de la santé et la recherche médicale (INSERM), Université Paris-Saclay (France), Paris Ile-de-France Region, and a PhD fellowship from Ministère de l’Enseignement Supérieur et de la Recherche (France) to A.S. We thank Dr. Katsuki Motoya, Prof. Sasaoka Toshikuni (Department of Comparative & Experimental Medicine/ Brain Research Institute; Niigata University, Japan), Dr. Jun Tanihata and Dr. Shin’ichi Takeda (National Center of Neurology and Psychiatry, Tokyo, Japan) for providing the *mdx52* mouse breeders. We are grateful to the Zootechnic platform of our institutes for mouse breeding and care.

## AUTHOR CONTRIBUTIONS

Conceptualization and Methodology. A.G.; Project design. funding acquisition and resources. A.G. and C.V.; Design of experiments. A.G., C.V. and O.V.; Production of AAV vectors. O.V.; Experimental work: O.V. A.S. M.D. X.P. Y.K. O.L.C. and C.N.; Data analysis. A.G. C.V. and O.V.; Original draft preparation. A.G. C.V. and O.V. All authors have read and agreed to the published version of the manuscript.

## DECLARATION OF INTERESTS

The authors declare no conflicts of interest.

## Notes

### Competing Interest Statement

The authors have declared no competing interest.

## REFERENCES

1. Muntoni, F., Torelli, S., and Ferlini, A. (2003). Dystrophin and mutations: one gene, several proteins, multiple phenotypes. Lancet Neurol 2, 731–740. 10.1016/s1474-4422(03)00585-4.

2. Hinton, V.J., Cyrulnik, S.E., Fee, R.J., Batchelder, A., Kiefel, J.M., Goldstein, E.M., Kaufmann, P., and De Vivo, D.C. (2009). Association of Autistic Spectrum Disorders With Dystrophinopathies. Pediatric Neurology 41, 339–346. 10.1016/j.pediatrneurol.2009.05.011.

3. Ricotti, V., Mandy, W.P.L., Scoto, M., Pane, M., Deconinck, N., Messina, S., Mercuri, E., Skuse, D.H., and Muntoni, F. (2016). Neurodevelopmental, emotional, and behavioural problems in Duchenne muscular dystrophy in relation to underlying dystrophin gene mutations. Developmental Medicine & Child Neurology 58, 77–84. 10.1111/dmcn.12922.

4. Colombo, P., Nobile, M., Tesei, A., Civati, F., Gandossini, S., Mani, E., Molteni, M., Bresolin, N., and D’Angelo, G. (2017). Assessing mental health in boys with Duchenne muscular dystrophy: Emotional, behavioural and neurodevelopmental profile in an Italian clinical sample. European Journal of Paediatric Neurology 21, 639–647. 10.1016/j.ejpn.2017.02.007.

5. Doorenweerd, N., Mahfouz, A., van Putten, M., Kaliyaperumal, R., T’ Hoen, P.A.C., Hendriksen, J.G.M., Aartsma-Rus, A.M., Verschuuren, J.J.G.M., Niks, E.H., Reinders, M.J.T., et al. (2017). Timing and localization of human dystrophin isoform expression provide insights into the cognitive phenotype of Duchenne muscular dystrophy. Sci Rep 7, 12575. 10.1038/s41598-017-12981-5.

6. Snow, W.M., Anderson, J.E., and Fry, M. (2014). Regional and genotypic differences in intrinsic electrophysiological properties of cerebellar Purkinje neurons from wild-type and dystrophin-deficient mdx mice. Neurobiol Learn Mem 107, 19–31. 10.1016/j.nlm.2013.10.017.

7. Hendriksen, J.G.M., and Vles, J.S.H. (2008). Neuropsychiatric disorders in males with duchenne muscular dystrophy: frequency rate of attention-deficit hyperactivity disorder (ADHD), autism spectrum disorder, and obsessive--compulsive disorder. J Child Neurol 23, 477–481. 10.1177/0883073807309775.

8. Vaillend, C., Aoki, Y., Mercuri, E., Hendriksen, J., Tetorou, K., Goyenvalle, A., and Muntoni, F. (2025). Duchenne muscular dystrophy: recent insights in brain related comorbidities. Nat Commun 16, 1298. 10.1038/s41467-025-56644-w.

9. Birnkrant, D.J., Bushby, K., Bann, C.M., Apkon, S.D., Blackwell, A., Brumbaugh, D., Case, L.E., Clemens, P.R., Hadjiyannakis, S., Pandya, S., et al. (2018). Diagnosis and management of Duchenne muscular dystrophy, part 1: diagnosis, and neuromuscular, rehabilitation, endocrine, and gastrointestinal and nutritional management. Lancet Neurol 17, 251–267. 10.1016/S1474-4422(18)30024-3.

10. Birnkrant, D.J., Bushby, K., Bann, C.M., Alman, B.A., Apkon, S.D., Blackwell, A., Case, L.E., Cripe, L., Hadjiyannakis, S., Olson, A.K., et al. (2018). Diagnosis and management of Duchenne muscular dystrophy, part 2: respiratory, cardiac, bone health, and orthopaedic management. The Lancet Neurology 17, 347–361. 10.1016/S1474-4422(18)30025-5.

11. Stahl, M., Zhu, Y., Goel, V., Leung, L., Carmack, T., Tami, Y., Hardin, T., Etxaniz, U., Kovach, P., Herzog, J., et al. AOC 1044 as a Novel Therapeutic Approach for DMD Patients Amenable to Exon 44 Skipping: EXPLORE44^TM^ Phase 1/2 Healthy Volunteer Data.

12. Barker, S.J., Thayer, M.B., Kim, C., Tatarakis, D., Simon, M.J., Dial, R., Nilewski, L., Wells, R.C., Zhou, Y., Afetian, M., et al. (2024). Targeting the transferrin receptor to transport antisense oligonucleotides across the mammalian blood-brain barrier. Sci Transl Med 16, eadi2245. 10.1126/scitranslmed.adi2245.

13. Aartsma-Rus, A., De Waele, L., Houwen-Opstal, S., Kirschner, J., Krom, Y.D., Mercuri, E., Niks, E.H., Straub, V., van Duyvenvoorde, H.A., and Vroom, E. (2023). The Dilemma of Choice for Duchenne Patients Eligible for Exon 51 Skipping The European Experience. J Neuromuscul Dis 10, 315–325. 10.3233/JND-221648.

14. Bardoni, A., Sironi, M., Felisari, G., Comi, G.P., and Bresolin, N. (1999). Absence of brain Dp140 isoform and cognitive impairment in Becker muscular dystrophy. Lancet 353, 897–898. 10.1016/S0140-6736(98)05801-2.

15. Giliberto, F., Ferreiro, V., Dalamon, V., and Szijan, I. (2004). Dystrophin deletions and cognitive impairment in Duchenne/Becker muscular dystrophy. Neurol Res 26, 83–87. 10.1179/016164104773026589.

16. Lidov, H.G.W., Selig, S., and Kunkel, L.M. (1995). Dp140: a novel 140 kDa CNS transcript from the dystrophin locus. Human Molecular Genetics 4, 329–335. 10.1093/hmg/4.3.329.

17. Saoudi, A., Zarrouki, F., Sebrié, C., Izabelle, C., Goyenvalle, A., and Vaillend, C. (2021). Emotional behavior and brain anatomy of the mdx52 mouse model of Duchenne muscular dystrophy. Dis Model Mech 14, dmm049028. 10.1242/dmm.049028.

18. Saoudi, A., Barberat, S., le Coz, O., Vacca, O., Doisy Caquant, M., Tensorer, T., Sliwinski, E., Garcia, L., Muntoni, F., Vaillend, C., et al. (2023). Partial restoration of brain dystrophin by tricyclo-DNA antisense oligonucleotides alleviates emotional deficits in mdx52 mice. Mol Ther Nucleic Acids 32, 173–188. 10.1016/j.omtn.2023.03.009.

19. Goyenvalle, A., Vulin, A., Fougerousse, F., Leturcq, F., Kaplan, J.-C., Garcia, L., and Danos, O. (2004). Rescue of dystrophic muscle through U7 snRNA-mediated exon skipping. Science 306, 1796–1799. 10.1126/science.1104297.

20. Vaillend, C., Perronnet, C., Ros, C., Gruszczynski, C., Goyenvalle, A., Laroche, S., Danos, O., Garcia, L., and Peltekian, E. (2010). Rescue of a dystrophin-like protein by exon skipping in vivo restores GABAA-receptor clustering in the hippocampus of the mdx mouse. Mol Ther 18, 1683–1688. 10.1038/mt.2010.134.

21. Dallérac, G., Perronnet, C., Chagneau, C., Leblanc-Veyrac, P., Samson-Desvignes, N., Peltekian, E., Danos, O., Garcia, L., Laroche, S., Billard, J.-M., et al. (2011). Rescue of a dystrophin-like protein by exon skipping normalizes synaptic plasticity in the hippocampus of the mdx mouse. Neurobiol Dis 43, 635–641. 10.1016/j.nbd.2011.05.012.

22. Goyenvalle, A., Babbs, A., van Ommen, G.-J.B., Garcia, L., and Davies, K.E. (2009). Enhanced exon-skipping induced by U7 snRNA carrying a splicing silencer sequence: Promising tool for DMD therapy. Mol Ther 17, 1234–1240. 10.1038/mt.2009.113.

23. Goyenvalle, A., and Davies, K.E. (2011). Engineering exon-skipping vectors expressing U7 snRNA constructs for Duchenne muscular dystrophy gene therapy. Methods Mol Biol 709, 179–196. 10.1007/978-1-61737-982-6_11.

24. Goyenvalle, A., Babbs, A., Wright, J., Wilkins, V., Powell, D., Garcia, L., and Davies, K.E. (2012). Rescue of severely affected dystrophin/utrophin-deficient mice through scAAV-U7snRNA-mediated exon skipping. Hum Mol Genet 21, 2559–2571. 10.1093/hmg/dds082.

25. Goyenvalle, A., Wright, J., Babbs, A., Wilkins, V., Garcia, L., and Davies, K.E. (2012). Engineering multiple U7snRNA constructs to induce single and multiexon-skipping for Duchenne muscular dystrophy. Mol Ther 20, 1212–1221. 10.1038/mt.2012.26.

26. Aupy, P., Zarrouki, F., Sandro, Q., Gastaldi, C., Buclez, P.-O., Mamchaoui, K., Garcia, L., Vaillend, C., and Goyenvalle, A. (2020). Long-Term Efficacy of AAV9-U7snRNA-Mediated Exon 51 Skipping in mdx52 Mice. Mol Ther Methods Clin Dev 17, 1037–1047. 10.1016/j.omtm.2020.04.025.

27. Hocquemiller, M., Giersch, L., Audrain, M., Parker, S., and Cartier, N. (2016). Adeno-Associated Virus-Based Gene Therapy for CNS Diseases. Hum Gene Ther 27, 478–496. 10.1089/hum.2016.087.

28. Kang, L., Jin, S., Wang, J., Lv, Z., Xin, C., Tan, C., Zhao, M., Wang, L., and Liu, J. (2023). AAV vectors applied to the treatment of CNS disorders: Clinical status and challenges. J Control Release 355, 458–473. 10.1016/j.jconrel.2023.01.067.

29. Lykken, E.A., Shyng, C., Edwards, R.J., Rozenberg, A., and Gray, S.J. (2018). Recent progress and considerations for AAV gene therapies targeting the central nervous system. Journal of Neurodevelopmental Disorders 10, 16. 10.1186/s11689-018-9234-0.

30. Cearley, C.N., and Wolfe, J.H. (2006). Transduction characteristics of adeno-associated virus vectors expressing cap serotypes 7, 8, 9, and Rh10 in the mouse brain. Molecular Therapy 13, 528–537. 10.1016/j.ymthe.2005.11.015.

31. Lidov, H.G.W., Byers, T.J., Watkins, S.C., and Kunkel, L.M. (1990). Localization of dystrophin to postsynaptic regions of central nervous system cortical neurons. Nature 348, 725–728. 10.1038/348725a0.

32. Holder, E., Maeda, M., and Bies, R.D. (1996). Expression and regulation of the dystrophin Purkinje promoter in human skeletal muscle, heart, and brain. Hum Genet 97, 232–239. 10.1007/BF02265272.

33. Fujimoto, T., Stam, K., Yaoi, T., Nakano, K., Arai, T., Okamura, T., and Itoh, K. (2023). Dystrophin Short Product, Dp71, Interacts with AQP4 and Kir4.1 Channels in the Mouse Cerebellar Glial Cells in Contrast to Dp427 at Inhibitory Postsynapses in the Purkinje Neurons. Mol Neurobiol 60, 3664–3677. 10.1007/s12035-023-03296-w.

34. Bremmer-Bout, M., Aartsma-Rus, A., de Meijer, E.J., Kaman, W.E., Janson, A.A.M., Vossen, R.H.A.M., van Ommen, G.-J.B., den Dunnen, J.T., and van Deutekom, J.C.T. (2004). Targeted exon skipping in transgenic hDMD mice: A model for direct preclinical screening of human-specific antisense oligonucleotides. Mol Ther 10, 232–240. 10.1016/j.ymthe.2004.05.031.

35. ’t Hoen, P.A.C., de Meijer, E.J., Boer, J.M., Vossen, R.H.A.M., Turk, R., Maatman, R.G.H.J., Davies, K.E., van Ommen, G.-J.B., van Deutekom, J.C.T., and den Dunnen, J.T. (2008). Generation and characterization of transgenic mice with the full-length human DMD gene. J Biol Chem 283, 5899– 5907. 10.1074/jbc.M709410200.

36. Araki, E., Nakamura, K., Nakao, K., Kameya, S., Kobayashi, O., Nonaka, I., Kobayashi, T., and Katsuki, M. (1997). Targeted Disruption of Exon 52 in the Mouse Dystrophin Gene Induced Muscle Degeneration Similar to That Observed in Duchenne Muscular Dystrophy. Biochemical and Biophysical Research Communications 238, 492–497. 10.1006/bbrc.1997.7328.

37. Bizot, F., Goossens, R., Tensorer, T., Dmitriev, S., Garcia, L., Aartsma-Rus, A., Spitali, P., and Goyenvalle, A. (2022). Histone deacetylase inhibitors improve antisense-mediated exon-skipping efficacy in *mdx* mice. Molecular Therapy - Nucleic Acids 30, 606–620. 10.1016/j.omtn.2022.11.017.

38. García-Rodríguez, R., Hiller, M., Jiménez-Gracia, L., van der Pal, Z., Balog, J., Adamzek, K., Aartsma-Rus, A., and Spitali, P. (2020). Premature termination codons in the DMD gene cause reduced local mRNA synthesis. Proceedings of the National Academy of Sciences 117, 16456–16464. 10.1073/pnas.1910456117.

39. Spitali, P., van den Bergen, J.C., Verhaart, I.E.C., Wokke, B., Janson, A.A.M., van den Eijnde, R., den Dunnen, J.T., Laros, J.F.J., Verschuuren, J.J.G.M., Hoen, P.A.C. ’t, et al. (2013). DMD transcript imbalance determines dystrophin levels. The FASEB Journal 27, 4909–4916. 10.1096/fj.13-232025.

40. Zhang, H., Yang, B., Mu, X., Ahmed, S.S., Su, Q., He, R., Wang, H., Mueller, C., Sena-Esteves, M., Brown, R., et al. (2011). Several rAAV vectors efficiently cross the blood-brain barrier and transduce neurons and astrocytes in the neonatal mouse central nervous system. Mol Ther 19, 1440–1448. 10.1038/mt.2011.98.

41. Hu, C., Busuttil, R.W., and Lipshutz, G.S. (2010). RH10 provides superior transgene expression in mice when compared with natural AAV serotypes for neonatal gene therapy. J Gene Med 12, 766–778. 10.1002/jgm.1496.

42. Zarrouki, F., Relizani, K., Bizot, F., Tensorer, T., Garcia, L., Vaillend, C., and Goyenvalle, A. (2022). Partial Restoration of Brain Dystrophin and Behavioral Deficits by Exon Skipping in the Muscular Dystrophy X-Linked (mdx) Mouse. Ann Neurol 92, 213–229. 10.1002/ana.26409.

43. Doisy, M., Vacca, O., Fergus, C., Gileadi, T., Verhaeg, M., Saoudi, A., Tensorer, T., Garcia, L., Kelly, V.P., Montanaro, F., et al. (2023). Networking to Optimize Dmd exon 53 Skipping in the Brain of mdx52 Mouse Model. Biomedicines 11, 3243. 10.3390/biomedicines11123243.

44. Saoudi, A., Fergus, C., Gileadi, T., Montanaro, F., Morgan, J.E., Kelly, V.P., Tensorer, T., Garcia, L., Vaillend, C., Muntoni, F., et al. (2023). Investigating the Impact of Delivery Routes for Exon Skipping Therapies in the CNS of DMD Mouse Models. Cells 12, 908. 10.3390/cells12060908.

45. Huda, F., Konno, A., Matsuzaki, Y., Goenawan, H., Miyake, K., Shimada, T., and Hirai, H. (2014). Distinct transduction profiles in the CNS via three injection routes of AAV9 and the application to generation of a neurodegenerative mouse model. Mol Ther Methods Clin Dev 1, 14032. 10.1038/mtm.2014.32.

46. Lee, N.K., Na, D.L., Kim, J.-W., Lee, B., Kim, H.J., Jang, H., Lee, J., and Chang, J.W. (2025). Evaluation of AAV transduction efficiency via multiple delivery routes: Insights from peripheral and central nervous system analysis. Neuroscience 573, 96–103. 10.1016/j.neuroscience.2025.03.026.

47. Naso, M.F., Tomkowicz, B., Perry, W.L., and Strohl, W.R. (2017). Adeno-Associated Virus (AAV) as a Vector for Gene Therapy. BioDrugs 31, 317–334. 10.1007/s40259-017-0234-5.

48. Wang, S., and Xiao, L. (2025). Progress in AAV-Mediated In Vivo Gene Therapy and Its Applications in Central Nervous System Diseases. International Journal of Molecular Sciences 26, 2213. 10.3390/ijms26052213.

49. Bey, K., Deniaud, J., Dubreil, L., Joussemet, B., Cristini, J., Ciron, C., Hordeaux, J., Le Boulc’h, M., Marche, K., Maquigneau, M., et al. (2020). Intra-CSF AAV9 and AAVrh10 Administration in Nonhuman Primates: Promising Routes and Vectors for Which Neurological Diseases? Molecular Therapy - Methods & Clinical Development 17, 771–784. 10.1016/j.omtm.2020.04.001.

50. D’Souza, P., Farmer, C., Johnston, J., Han, S.T., Adams, D., Hartman, A.L., Zein, W., Huryn, L.A., Solomon, B., King, K., et al. (2024). GM1 Gangliosidosis Type II: Results of a 10-Year Prospective Study. medRxiv, 2024.01.04.24300778. 10.1101/2024.01.04.24300778.

51. Nagy, A., Bley, A.E., and Eichler, F. (1993). Canavan Disease. In GeneReviews®, M. P. Adam, J. Feldman, G. M. Mirzaa, R. A. Pagon, S. E. Wallace, and A. Amemiya, eds. (University of Washington, Seattle).

52. Gao, C., Pielas, M., Jiao, F., Mei, D., Wang, X., Kotulska, K., and Jozwiak, S. (2023). Epilepsy in Dravet Syndrome-Current and Future Therapeutic Opportunities. J Clin Med 12, 2532. 10.3390/jcm12072532.

53. Hammond, S.L., Leek, A.N., Richman, E.H., and Tjalkens, R.B. (2017). Cellular selectivity of AAV serotypes for gene delivery in neurons and astrocytes by neonatal intracerebroventricular injection. PLoS One 12, e0188830. 10.1371/journal.pone.0188830.

54. Benkhelifa-Ziyyat, S., Besse, A., Roda, M., Duque, S., Astord, S., Carcenac, R., Marais, T., and Barkats, M. (2013). Intramuscular scAAV9-SMN injection mediates widespread gene delivery to the spinal cord and decreases disease severity in SMA mice. Mol Ther 21, 282–290. 10.1038/mt.2012.261.

55. Grimm, D., Lee, J.S., Wang, L., Desai, T., Akache, B., Storm, T.A., and Kay, M.A. (2008). In vitro and in vivo gene therapy vector evolution via multispecies interbreeding and retargeting of adeno-associated viruses. J Virol 82, 5887–5911. 10.1128/JVI.00254-08.

56. Koerber, J.T., Klimczak, R., Jang, J.-H., Dalkara, D., Flannery, J.G., and Schaffer, D.V. (2009). Molecular evolution of adeno-associated virus for enhanced glial gene delivery. Mol Ther 17, 2088–2095. 10.1038/mt.2009.184.

57. Deverman, B.E., Pravdo, P.L., Simpson, B.P., Kumar, S.R., Chan, K.Y., Banerjee, A., Wu, W.-L., Yang, B., Huber, N., Pasca, S.P., et al. (2016). Cre-dependent selection yields AAV variants for widespread gene transfer to the adult brain. Nat Biotechnol 34, 204–209. 10.1038/nbt.3440.

58. Chauhan, M., Daugherty, A.L., Khadir, F. (Ellie), Duzenli, O.F., Hoffman, A., Tinklenberg, J.A., Kang, P.B., Aslanidi, G., and Pacak, C.A. (2024). AAV-DJ is superior to AAV9 for targeting brain and spinal cord, and de-targeting liver across multiple delivery routes in mice. Journal of Translational Medicine 22, 824. 10.1186/s12967-024-05599-5.

59. Chan, K.Y., Jang, M.J., Yoo, B.B., Greenbaum, A., Ravi, N., Wu, W.-L., Sánchez-Guardado, L., Lois, C., Mazmanian, S.K., Deverman, B.E., et al. (2017). Engineered AAVs for efficient noninvasive gene delivery to the central and peripheral nervous systems. Nat Neurosci 20, 1172–1179. 10.1038/nn.4593.

60. Gharibi, S., Vaillend, C., and Lindsay, A. (2024). The unconditioned fear response in vertebrates deficient in dystrophin. Progress in Neurobiology 235, 102590. 10.1016/j.pneurobio.2024.102590.

61. Goyenvalle, A., Vulin, A., Fougerousse, F., Leturcq, F., Kaplan, J.-C., Garcia, L., and Danos, O. (2004). Rescue of dystrophic muscle through U7 snRNA-mediated exon skipping. Science 306, 1796–1799. 10.1126/science.1104297.

62. McCarty, D.M., Monahan, P.E., and Samulski, R.J. (2001). Self-complementary recombinant adeno-associated virus (scAAV) vectors promote efficient transduction independently of DNA synthesis. Gene Ther 8, 1248–1254. 10.1038/sj.gt.3301514.

63. Fu, H., Muenzer, J., Samulski, R.J., Breese, G., Sifford, J., Zeng, X., and McCarty, D.M. (2003). Self-complementary adeno-associated virus serotype 2 vector: global distribution and broad dispersion of AAV-mediated transgene expression in mouse brain. Mol Ther 8, 911–917. 10.1016/j.ymthe.2003.08.021.

64. Hironaka, K., Yamazaki, Y., Hirai, Y., Yamamoto, M., Miyake, N., Miyake, K., Okada, T., Morita, A., and Shimada, T. (2015). Enzyme replacement in the CSF to treat metachromatic leukodystrophy in mouse model using single intracerebroventricular injection of self-complementary AAV1 vector. Sci Rep 5, 13104. 10.1038/srep13104.

65. Dias Florencio, G., Precigout, G., Beley, C., Buclez, P.-O., Garcia, L., and Benchaouir, R. (2015). Simple downstream process based on detergent treatment improves yield and *in vivo* transduction efficacy of adeno-associated virus vectors. Molecular Therapy - Methods & Clinical Development 2, 15024. 10.1038/mtm.2015.24.

66. Paylor, R., Zhao, Y., Libbey, M., Westphal, H., and Crawley, J.N. (2001). Learning impairments and motor dysfunctions in adult *Lhx5*-deficient mice displaying hippocampal disorganization. Physiology & Behavior 73, 781–792. 10.1016/S0031-9384(01)00515-7.

67. Paylor, R., Zhao, Y., Libbey, M., Westphal, H., and Crawley, J.N. (2001). Learning impairments and motor dysfunctions in adult *Lhx5*-deficient mice displaying hippocampal disorganization. Physiology & Behavior 73, 781–792. 10.1016/S0031-9384(01)00515-7.

68. Vaillend, C., and Chaussenot, R. (2017). Relationships linking emotional, motor, cognitive and GABAergic dysfunctions in dystrophin-deficient mdx mice. Human Molecular Genetics 26, 1041–1055. 10.1093/hmg/ddx013.

