## Supplemental informations for "Ineffective behavioral rescue despite partial brain Dp427 restoration by AAV9-U7-mediated exon 51 skipping in *mdx52* mice"

### SUPPLEMENTAL INFORMATION

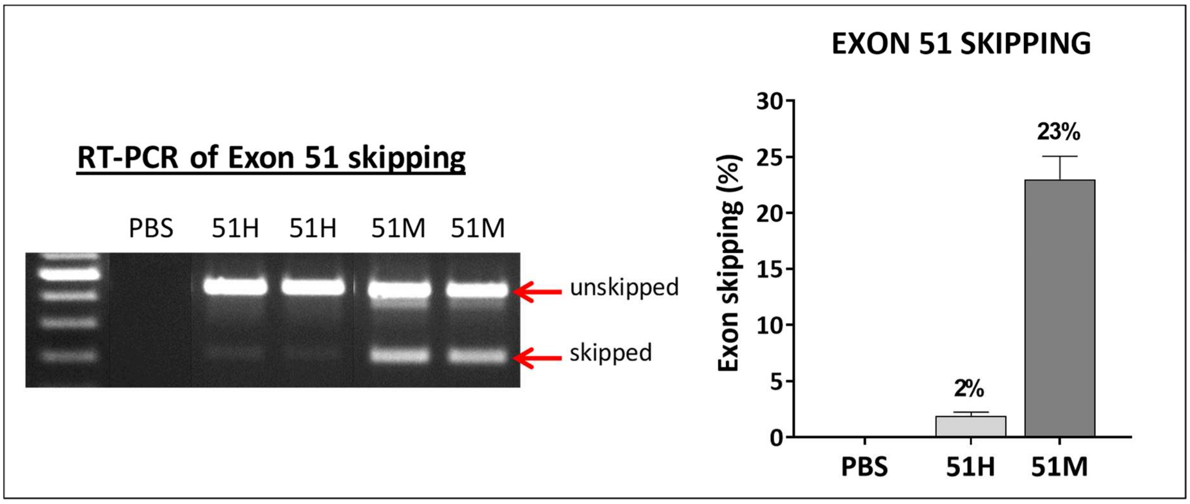


<u>Figure S1</u>: **Comparison of exon 51 skipping efficiency using U7-exon51 with human versus murine sequences in *mdx52* mice.** (Left panel): RT-PCR analysis of exon 51 skipping after intramuscular injection of scAAV9-U7-51H (human sequence) or scAAV9-U7-51M (murine sequence) in *mdx52* mice at an equivalent dose of $1\times10^{12}$ vg. Lane 1: DNA ladder; Lane 2: PBS-injected *mdx52* mouse (negative control); Lanes 3 & 4: scAAV9-U7-51H-injected *mdx52* mice; Lanes 5 & 6: scAAV9-U7-51M-injected *mdx52* mice. (Right panel): Quantification of exon 51 skipping efficiency as a percentage, based on RT-PCR gel analysis. Data are presented as mean ± SEM and analyzed using Unpaired t test test; ***$p < 0.001$.

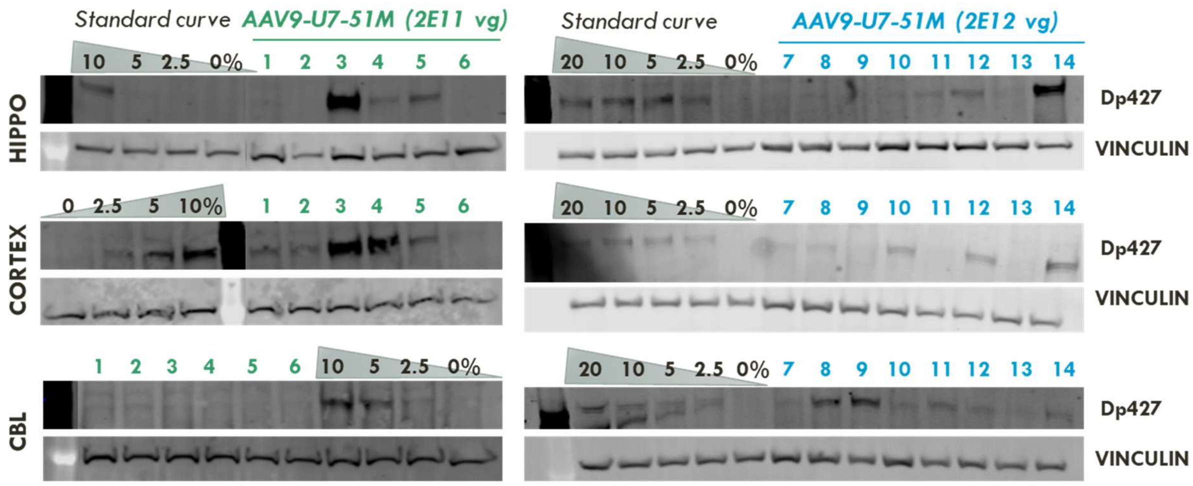


**Figure S2: Western blot analysis of Dp427 restoration in *mdx52* mouse brain after AAV9-U7-51M administration.** Representative western blot images of Dp427 and vinculin (loading control) in hippocampus (HIPPO), cortex, and cerebellum (CBL) from *mdx52* mice injected intracerebroventricularly with AAV9-U7-51M at two doses: 2E11 vg (mice 1–6, green) and 2E12 vg (mice 7–14, blue). A standard curve was loaded alongside, corresponding to protein extracts from wild-type mouse brain diluted into *mdx52* brain extracts to yield 20%, 10%, 5%, 2.5% and 0% of normal Dp427 levels, as indicated. Restoration of Dp427 was variable across animals and brain regions.

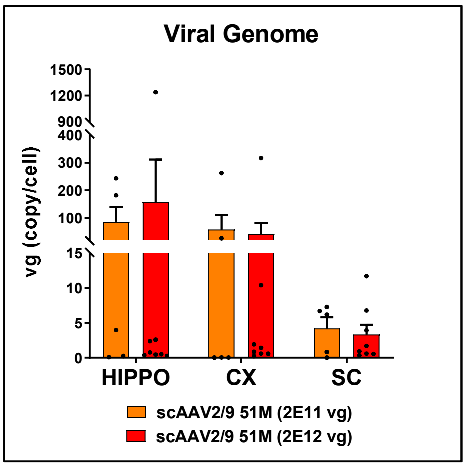


**Figure S3: Biodistribution of the scAAV-U7-51M vector.** Quantification of the viral genome copies via qPCR on genomic DNA in the HIP, CX and SC, 9 weeks after scAAV-U7-51M ICV injection, comparing low- and high-dose treatments (n=7 for the low-dose in orange; n=8 for the high dose in red).

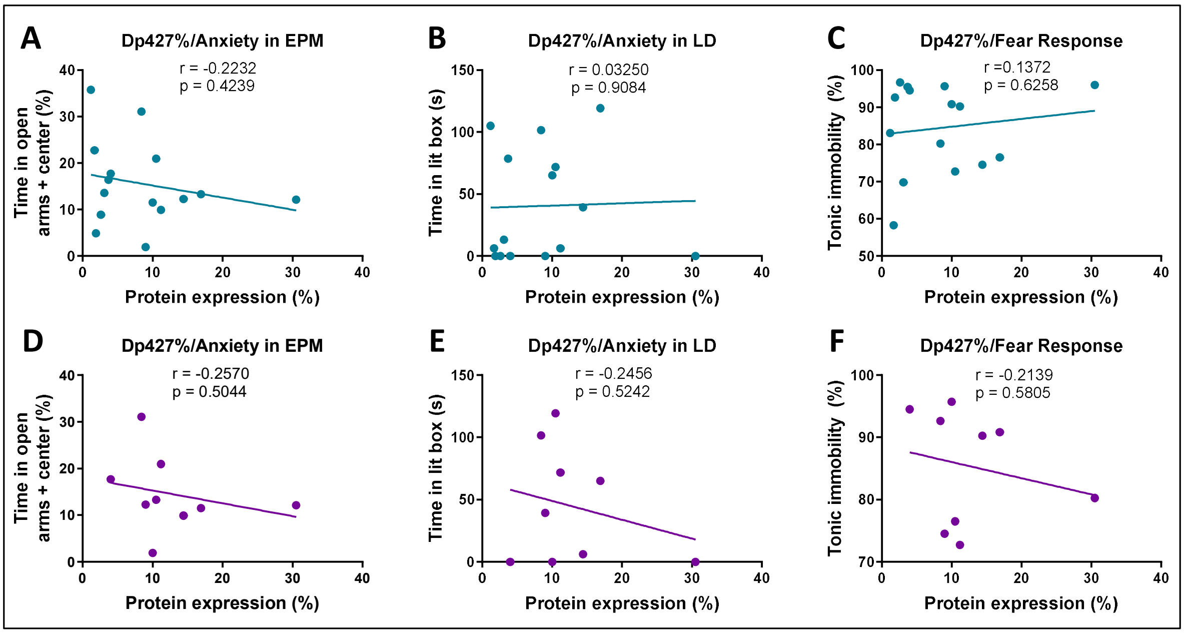


**Figure S4: Correlation of Dp427 Restoration with Anxiety- and Fear-Related Behaviors.** This figure displays the correlation between the percentage of Dp427 restoration and behavioral measures of anxiety and fear in treated *mdx52* mice. Panels A–C show correlations for all mice (n=15) in the study, while panels D–F present correlations for the post-hoc selected subset of mice (n=9). Specific behavioral parameters are: time spent in open arms and in the center in the elevated plus maze (EPM) (A, D), time spent in the lit box in the light-dark choice test (LD) (B, E), and tonic immobility in the unconditioned fear response test (C, F). Pearson correlation coefficients (r), P values (p) and linear regression lines are shown.
